# Pan-cancer profiling of tumor-infiltrating natural killer cells through transcriptional reference mapping

**DOI:** 10.1101/2023.10.26.564050

**Authors:** Herman Netskar, Aline Pfefferle, Jodie P Goodridge, Ebba Sohlberg, Olli Dufva, Sara A Teichmann, Trevor Clancy, Amir Horowitz, Karl-Johan Malmberg

## Abstract

The functional diversity of natural killer (NK) cell repertoires stems from differentiation, homeostatic receptor-ligand interactions, and adaptive-like responses to viral infections. Here, we generated a single-cell transcriptional reference map of healthy human blood and tissue-derived NK cells, with temporal resolution and fate-specific expression of gene regulator networks defining NK cell differentiation. Using transfer learning, transcriptomes of tumor-infiltrating NK cells from seven solid tumor types (427 patients), combined from 39 datasets, were incorporated into the reference map and interrogated for tumor microenvironment (TME)-induced perturbations. We identified six functionally distinct NK cellular states in healthy and malignant tissues, two of which were commonly enriched for across tumor types: a dysfunctional ‘stressed’ CD56^bright^ state susceptible to TME-induced immunosuppression and a cytotoxic TME-resistant ‘effector’ CD56^dim^ state. The ratio of ‘stressed’ CD56^bright^ and ‘effector’ CD56^dim^ was predictive of patient outcome in malignant melanoma and osteosarcoma. This resource may inform the design of novel NK cell therapies and can be extended endlessly through transfer learning to interrogate new datasets from experimental perturbations or disease conditions.

## Introduction

Natural killer (NK) cells are innate lymphocytes that play a vital role in the immune response through their ability to directly kill transformed and virus infected cells, and by orchestrating the early phase of the adaptive immune response^1^. NK cells are commonly divided into two functionally distinct subsets, CD56^bright^ and CD56^dim^ NK cells^2, 3^. However, this is an oversimplified view of the repertoire. Mass cytometry profiling of NK cell repertoires at the single cell level revealed an extensive phenotypic diversity comprising up to 100,000 unique subsets in healthy individuals^4^. Much of this diversity is based on combinatorial expression of stochastically expressed germline encoded activating and inhibitory receptors that bind to HLA class I and tune NK cell function in a process termed NK cell education^5, 6^. Another layer of diversity reflects the continuous differentiation through well-defined intermediate phenotypes from the naïve CD56^bright^ NK cells through CD62L^+^NKG2A^+^KIR^-^CD57^-^CD56^dim^ NK cells to terminally differentiated, adaptive CD62L^-^NKG2C^+^CD57^+^KIR^+^CD56^dim^ NK cells, associated with past infection of cytomegalovirus (CMV)^7, 8, 9, 10^. Given the increasing interest to harness the cytolytic potential of NK cells in cell therapy against cancer, it is of fundamental importance to understand the molecular programs and gene regulatory circuits driving NK cell differentiation and the underlying functional diversification of the human NK cell repertoire.

Utilizing single-cell RNA sequencing (scRNA-seq), Crinier et al. discovered organ-specific signatures in human spleen NK cells and two major transcriptional clusters in blood-derived NK cells (PB-NK), corresponding to CD56^dim^ (NK1) and CD56^bright^ (NK2) NK cell subsets^2^. Bulk RNA and ChIP sequencing identified dominant transcription factor (TF) axes defining CD56^bright^ (TCF1-LEF-MYC) and CD56^dim^ (PRDM1) phenotypic subsets, respectively^11^. Later research reported additional diversity with unique transcriptional clusters, including IL-2 and type I interferon-responding NK cell subsets^12^ and an intermediate CD56^dim^GzmK^+^ stage, potentially bridging CD56^bright^ and CD56^dim^ NK cells^13^. A comprehensive analysis unveiled Bcl11b’s role in driving NK cell differentiation towards the adaptive state, reciprocally suppressing early TFs like RUNX2 and ZBTB16^14^. Combining gene expression analysis, chromatin accessibility, and lineage tracing via mitochondrial DNA (mtDNA) mutations, Rückert et al. revealed a clonal expansion and a distinct inflammatory memory signature in adaptive NK cells^15^. Using a pan-cancer single-cell atlas approach, Tang et al.^16^ identified a tumor-enriched dysfunctional CD56^dim^ CD16^hi^ NK cell population interacting with LAMP3^+^ dendritic cells in the tumor microenvironment (TME). Hence, scRNA-seq and bulk RNA-seq usage has defined major transcriptional regulatory hubs during NK cell differentiation and identified a persistent memory state in human innate immunity. However, it remains unclear how the regulatory gene circuits that operate under homeostasis in healthy tissues are affected by cellular and/or soluble cues in the TME, resulting in perturbed functional states within tumor-infiltrating NK cells.

Here we establish a single-cell transcriptional reference map that resolves gene expression trends and dominating TF-target interactions during NK cell differentiation in blood and normal tissues. Reference mapping enables the analysis of cellular differences and gene programs in diseases and various conditions by contextualizing new datasets within a healthy transcriptional reference, facilitating the identification of novel states not found in the reference^17^. We utilize our NK cell reference map, compiled from 44,640 PB-NK cells (12 donors) and 27,489 tissue resident NK (TrNK) cells (136 donors), to query the regulones and functional states, as defined through gene expression signatures, of tumor infiltrating NK (TiNK) cells derived from 427 patients with seven distinct solid tumors (38,862 TiNKs). We found that TrNK and TiNK cells have a clear tissue residency signature but still share the dominant regulons of blood CD56^bright^ and CD56^dim^ NK cells. Of the six functional states identified in our pan-cancer atlas, a dysfunctional ‘stressed’ CD56^bright^ state susceptible to TME-associated cellular communication and a cytotoxic ‘effector’ CD56^dim^ state resistant to TME-associated cellular communication were commonly enriched across tumor types. Stratification of patient survival data identified a high ratio of ‘effector’ CD56^dim^ to ‘stressed’ CD56^bright^ state to correlate with improved survival in osteosarcoma and melanoma patients. This resource provides a granular view of cancer-specific alterations of solid-tumor infiltrating NK cells, identifying how the TME can lead to NK cell dysfunction and may inspire new strategies to engineer cell therapy products with robust functional phenotypes resistant to TME-induced suppressive mechanisms.

## Results

### NK cell subset annotation of single-cell RNA sequencing data using predictive gene signatures

To establish a pan-cancer atlas of tumor-infiltrating NK cells, we first defined NK cell differentiation at the transcriptional level. We performed single-cell RNA sequencing (scRNA-seq) of the total NK cell population from 7 healthy donors and integrated our transcriptomes with 5 publicly available donor datasets^2, 18^ using scVI^19^ (**Supplemental Table 1**). By retaining only cell-to-cell variation independent from sample-to-sample variation, the initial clustering by donor and laboratory origin was successfully integrated into a homogenous population of cells and visualized using diffusion maps^20^ to preserve the continuous trajectories observed with biological differentiation (**Figure 1A**). Although NK differentiation is best described as a continuum, CD56^bright^ and CD56^dim^ NK cell represent two distinct stages of differentiation. By performing gene signature scoring using AUCell^21^, we identified cells at the top of the diffusion map embedding scoring high for the CD56^bright^ gene signature^2^, while the main body of the embedding exhibited increasing intensity of the CD56^dim^ signature^2^ (**Figure 1B**). Scoring of two independent gene signatures based on the CD56^bright/dim^ regulon^11^ and proteome^22^ confirmed our results (**Supplemental Figure 1A-B**).

**Figure 1.**
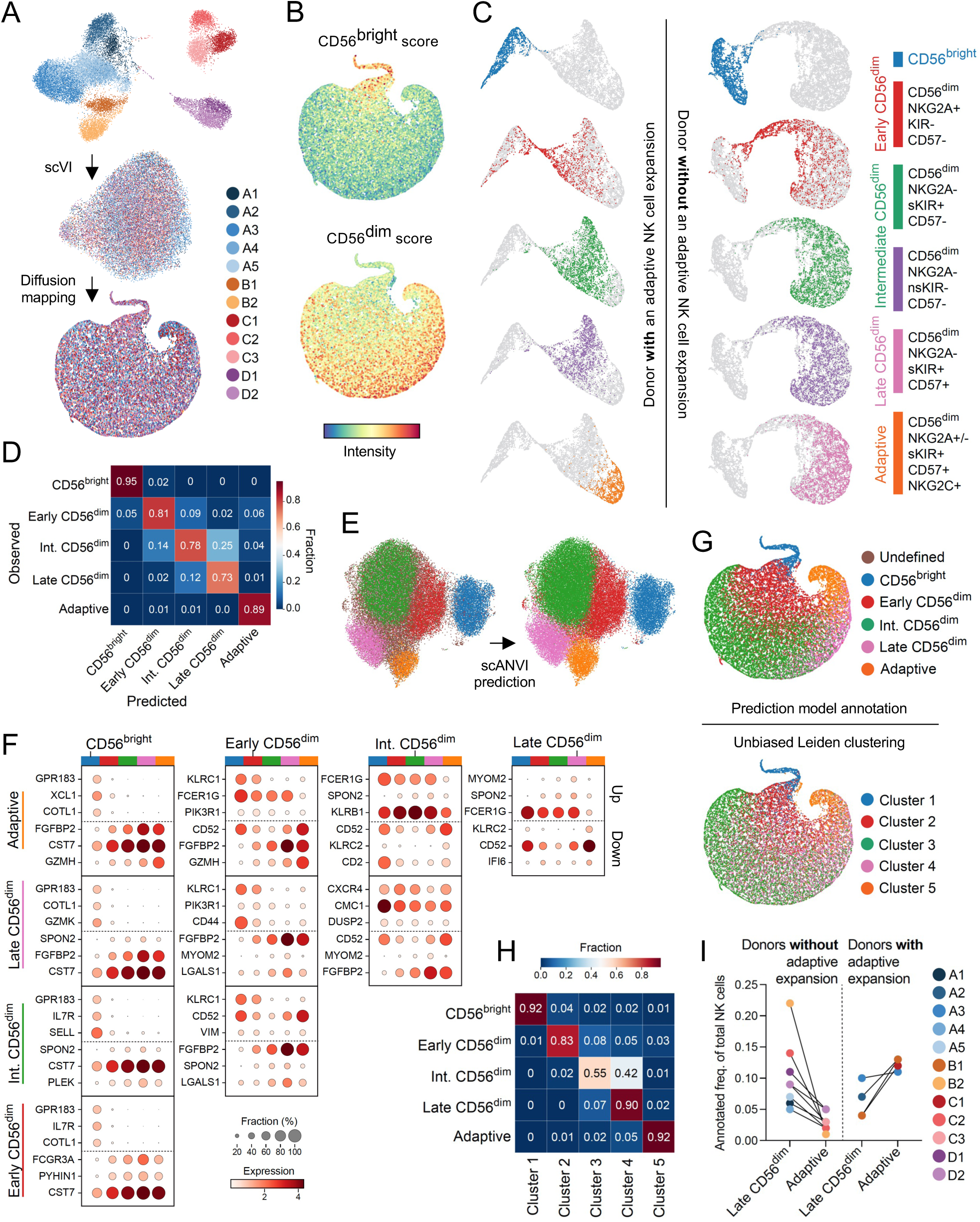
NK cell differentiation at the transcriptional level. (**A**) Integration process of scRNA- seq data of NK cells from 12 donors and 4 different laboratories using scVI showing UMAP representation followed by diffusion mapping. (**B**) AUCell scores of gene signatures for CD56^bright^ and CD56^dim^ NK cell subsets. (**C**) UMAP representation of 5 sorted subsets from a donor with an adaptive expansion (left) and a donor without an adaptive expansion (right). (**D**) Heatmap depicting accuracy of our prediction model for subset annotation tested on 15% of heldout cells from the subset-specific dataset (2 donors). (**E**) UMAP representation showing annotation of NK cells (12 donors, left) with subset labels (right) using a scANVI model trained with sorted subset data (2 donors) and the final diffusion map depicting subset annotations. (**F**) Dotplots showing the top three up and downregulated genes between all pairs of subsets as identified by the differential expression module in scANVI, visualized across the differentiation spectrum. (**G**) Diffusion map depicting Leiden clustering of the 12 donor NK cell dataset. (**H**) Heatmap showing distribution of our annotated 12 donor NK cell subsets over the 5 Leiden clusters. (**I**) Frequency of annotated late CD56^dim^ and adaptive NK cell subsets in donors with and without an adaptive NK cell expansion.

The relatively large and heterogeneous population of CD56^dim^ NK cells is commonly phenotypically defined into functionally distinct subsets based on a selected number of inhibitory and activating receptors contributing to the functional tuning^7^. To identify predictive gene signatures associated with these functional stages encompassing NK cell differentiation, we sorted and sequenced equal numbers of CD56^bright^ NK cells and four CD56^dim^ NK cell subsets (NKG2A^+^KIR^-^CD57^-^, NKG2A^-^self-KIR^+^CD57^-^, NKG2A^-^nonself-KIR^+^CD57^-^, NKG2A^-^self-KIR^+^CD57^+^ or NKG2A^-/+^self-KIR^+^CD57^+^NKG2C^+^) from two donors, one without and one with a large adaptive NK cell expansion (**Figure 1C**, **Supplemental Figure 1C-D**). Transcriptionally, the adaptive NK cell subset was the most distinct as the remaining CD56^dim^ subsets exhibited a high degree of transcriptional overlap, while still ordering themselves along the previously defined maturation scheme (**Figure 1C**). As previously observed in bulk RNA-seq data^23^, the transcriptomes of self and non-self KIR^+^ NK cells were highly similar even at the single cell level and thus merged for subsequent analysis (**Figure 1C**). The five transcriptionally distinct NK subsets were renamed to reflect their maturation stage: ‘CD56^bright^’, ‘early CD56^dim^’, ‘intermediate CD56^dim^’, ‘late CD56^dim^’ and ‘adaptive’ (**Figure 1C**).

We next utilized a semi-supervised model, scANVI^24^, to leverage our identified NK cell subset gene signatures to predict and infer subset annotation of compiled bulk NK cell scRNA-seq datasets. We first tested the accuracy of the prediction model (M1) on 15% of the subset-sorted NK cells (**Figure 1C**) which were not included in the training of the model. Transcriptionally distinct subsets (CD56^bright^, adaptive) were annotated with high accuracy, while subsets exhibiting higher transcriptional overlap were annotated with slightly reduced accuracy (**Figure 1D**). Implementing the model, we could annotate the total NK cell dataset comprising 23,253 single cell transcriptomes (12 donors) at the subset level (**Figure 1E**). The models top three differentially expressed genes (up and down) for each subset’s gene signature showed some overlap, further highlighting the continuous nature of NK cell differentiation at the transcriptional level (**Figure 1F**). To validate our annotation model, we performed unbiased clustering (Leiden) of the total NK cell dataset (12 donors), identifying five clusters closely matching our annotated five NK cell subsets (**Figure 1G**). A small portion of intermediate CD56^dim^ annotated NK cells clustered together with late CD56^dim^ annotated NK cells in cluster 4 (**Figure 1H**), likely corresponding to more mature cells within the population. Having confirmed the validity of our 5 NK cell subsets, M1 was utilized to identify donors with an adaptive NK cell expansion, which were all confirmed to be CMV seropositive (**Figure 1I**). Thus, this first scANVI model forms a basis to interrogate cellular states layered on top of the natural transcriptional changes with NK cell subsets at different stages of differentiation.

### Temporal resolution of gene regulator networks with fate-specific expression

To decipher the regulatory gene pathways driving NK cell differentiation at the transcriptional level, we implemented two different methods to calculate pseudotime, namely Palantir^25^ and RNA velocity-based pseudotime^26, 27^. Palantir identifies terminal cells based on a chosen starting cell, placing the remaining cells along a timeline (pseudotime). Defining the starting cell (blue) based on the lowest CD56^dim^ score^2^ (**Figure 1B**) identified two terminal cells (orange), predicted to be part of the late CD56^dim^ and adaptive population respectively (**Figure 2A**). To validate this trajectory, we utilized the dynamic model implemented in scVelo^26^ to compute RNA velocity (spliced versus unspliced transcripts), inferring pseudotime without a predefined starting cell (**Supplemental Figure 2A- B**). The resulting vector field and extrapolated pseudotime confirmed a trajectory starting within the CD56^bright^ NK cell subset and terminating in the adaptive subset (**Figure 2B**). Lastly, to infer developmental relationships at the resolution of the five subsets, representing functionally distinct subsets and proposed stages of NK cell differentiation^7^, we applied Partition-based graph abstraction (PAGA)^28^ to quantify their connectivity and estimate transitions. In line with the two terminal fates (late CD56^dim^, adaptive) identified by Palantir, we analyzed conventional and adaptive donors separately (**Figure 1I**). In both types of donors, early CD56^dim^ NK cells formed the connecting link between CD56^bright^ and the remaining CD56^dim^ populations (**Figure 2C-D**). However, while adaptive donors NK cells continued their progression to intermediate CD56^dim^ cells, terminating in the adaptive population, conventional donors instead branched into the intermediate or late CD56^dim^ populations (**Figure 2C-D**).

**Figure 2.**
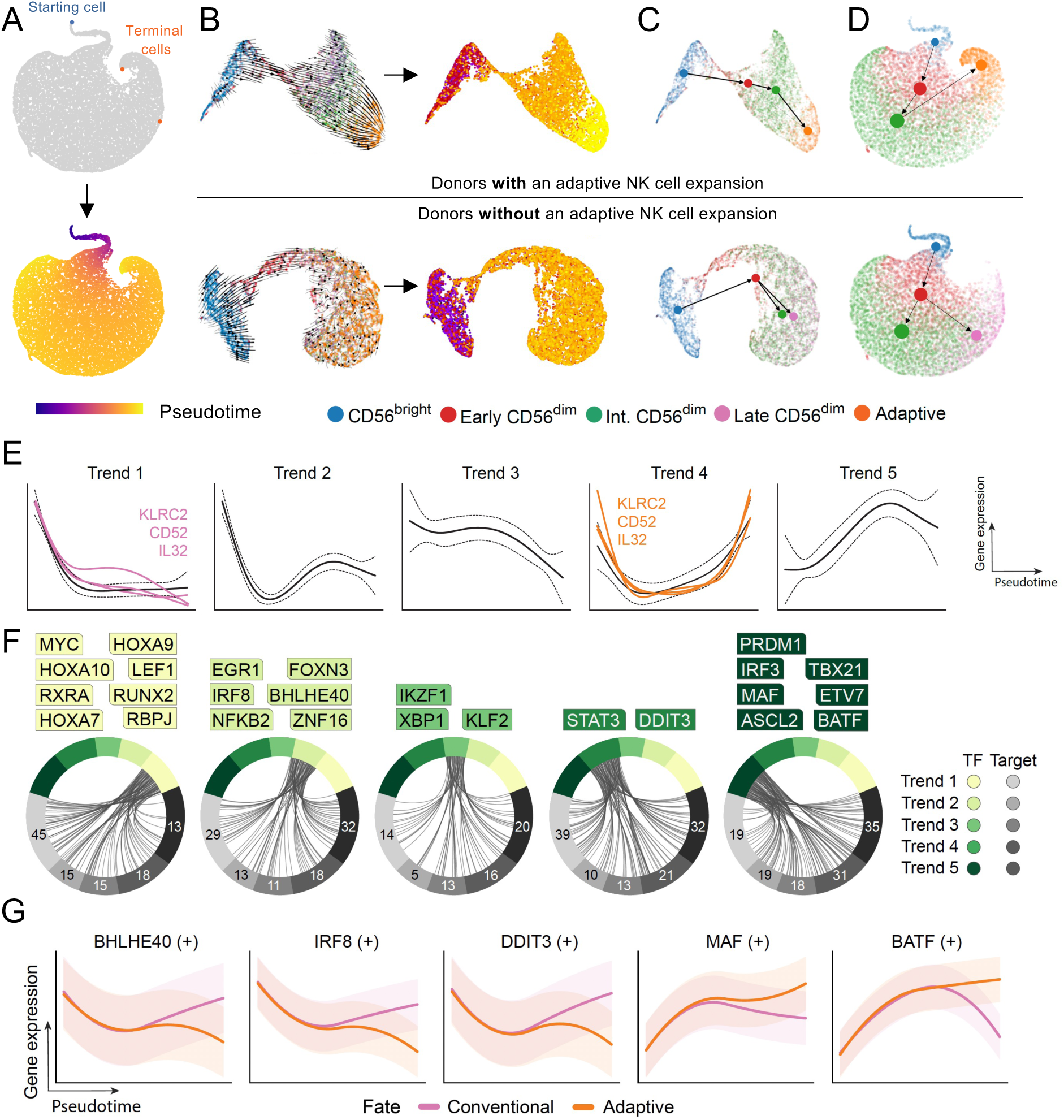
Gene regulatory networks defining conventional and adaptive NK cell fates. (**A**) UMAP representation highlighting the starting cell (blue) with the lowest value CD56^dim^ signature score and the two terminal cells (orange) as predicted by Palantir. (**B**) UMAP representation of the data from the sorted subsets (2 donors) showing the RNA velocity vector field as a stream plot and the inferred pseudotime. (**C-D**) PAGA graph with directionality and transitions from RNA velocity analysis for the sorted subsets (2 donors) (**C**) subset-inferred bulk donors (12 donors) stratified based on presence or absence of adaptive expansion (**D**). (**E**) Gene trends clustered into five overall trends of expression along pseudotime, showing expression of KLRC2, CD52 and IL32 in both terminal fates (pink = conventional fate, orange = adaptive fate). (**F**) Inferred gene regulatory networks where dominant transcription factors for each trend are highlighted. (**G**) Selection of regulons showing differential expression over pseudotime within the conventional and adaptive fate.

Having established a temporal axis to NK cell differentiation, we utilized generalized additive models to compute gene expression trends as a function of time for each gene^25^, which clustered into five distinct trends (**Figure 2E**). Genes varying in expression across the two terminal fates were depicted in their trends for each fate, exemplified by KLRC2, CD52^15, 18^, IL32 clustering into Trend 1 in the conventional late CD56^dim^ fate and into Trend 4 in the adaptive fate (**Figure 2E**). Based on the two-fate model, we constructed gene regulatory networks (GRN)^21^ stratified by the five gene trends, identifying the dominant TFs across pseudotime and their known downstream target genes (**Figure 2F**). Trend 1 is dominated by genes which are downregulated with differentiation from CD56^bright^ to CD56^dim^ cells, including previously reported TFs (MYC, LEF1, RUNX2)^11^, RBPJ^29^ involved in Notch signaling, the retinoic acid receptor (RXRA), and TFs regulating ID2 expression (HOXA9, HOXA10)^30^ (**Figure 2E-F**). Trend 2 genes, compared to Trend 1, are upregulated during differentiation from early to intermediate CD56^dim^ cells and include among others EGR1^31^ (cell survival, proliferation, apoptosis, regulates TRAIL expression), BHLHE40^32, 33^ (associated with NK cell activation and represses RXRA) and IRF8^34, 35^ (role in orchestrating adaptive response, essential NK cell gene) (**Figure 2E-F**). TFs exhibiting less dynamic changes across pseudotime are clustered in Trend 3, such as IKZF1, XBP1 and KLF2 which play a role in regulating homeostatic proliferation, effector function and cytokine responsiveness^36, 37^. TFs exhibiting higher expression at the start and end of pseudotime fall into Trend 4, including STAT3 (cell survival, IFNγ production) and DDIT3^38^ (stress response, metabolism). Lastly, expression of Trend 5 genes steadily increases with differentiation, decreasing only during late differentiation and includes previously reported TFs associated with CD56^dim^ NK cells (MAF, PRDM1, TBX21))^11^, the AP-1 family member BATF, the ETS family member ETV7, and the Wnt target gene ASCL2 (**Figure 2E-F**). The TF-based GRNs were further curated to only retain direct targets with significant motif enrichment, referred to as ‘regulons’ (denoted by ‘(+)’), expression of which was confirmed in an independent bulk RNA-seq data set on sorted NK cell subsets. Regulon expression substantially differing between the conventional and adaptive fate include conventional fate associated BHLHE40^33^, IRF8^34, 35^ and DDIT3^38^ and adaptive fate associated MAF^11^ and BATF regulons (**Figure 2G**). Clustering dominant TFs according to their temporal expression during NK cell differentiation revealed a set of highly connected regulatory circuits, expression of which diverged during terminal differentiation into one of the two cell fates, conventional or adaptive.

### Transfer learning to generate pan-cancer atlas of tissue-derived and solid tumor-infiltrating NK cells

Having transcriptionally defined NK cell differentiation in peripheral blood, we proceeded to train a second model (M2) with publicly available scRNA-seq datasets encompassing six healthy tissues (brain, breast, lung, pancreas, prostate, skin) from a total of 136 donors using scVI^19^ (**Figure 3A, Supplemental Table 2**). The tissue-specific datasets were integrated and annotated using scANVI, and CellTypist^39^ was used to identify immune subsets of interest (**Figure 3B, Supplemental Figure 3A-E**). The annotated tissue-derived CD56^bright^ and CD56^dim^ NK cell populations (27,489 cells) were extracted from the datasets and integrated into our reference map (**Figure 3C**). Tissue-residency status was confirmed by scoring for a tissue residency (Tr) signature, which was most pronounced in TrCD56^bright^ NK cells but also increased in TrCD56^dim^ NK cells (**Figure 3D**). CD56^bright^ and CD56^dim^ subsets from peripheral blood and tissues clustered together (**Figure 3E**) and were more tightly connected than to their respective tissues, apart from skin-derived NK cells (**Figure 3F**). Therefore, differentiation stage has a greater influence on the NK cell transcriptome compared to tissue origin. Notably, CellTypist did not identify a CD56^bright^ NK cell population in neither brain nor breast tissue (**Supplemental Figure 3A, Figure 3E**).

**Figure 3.**
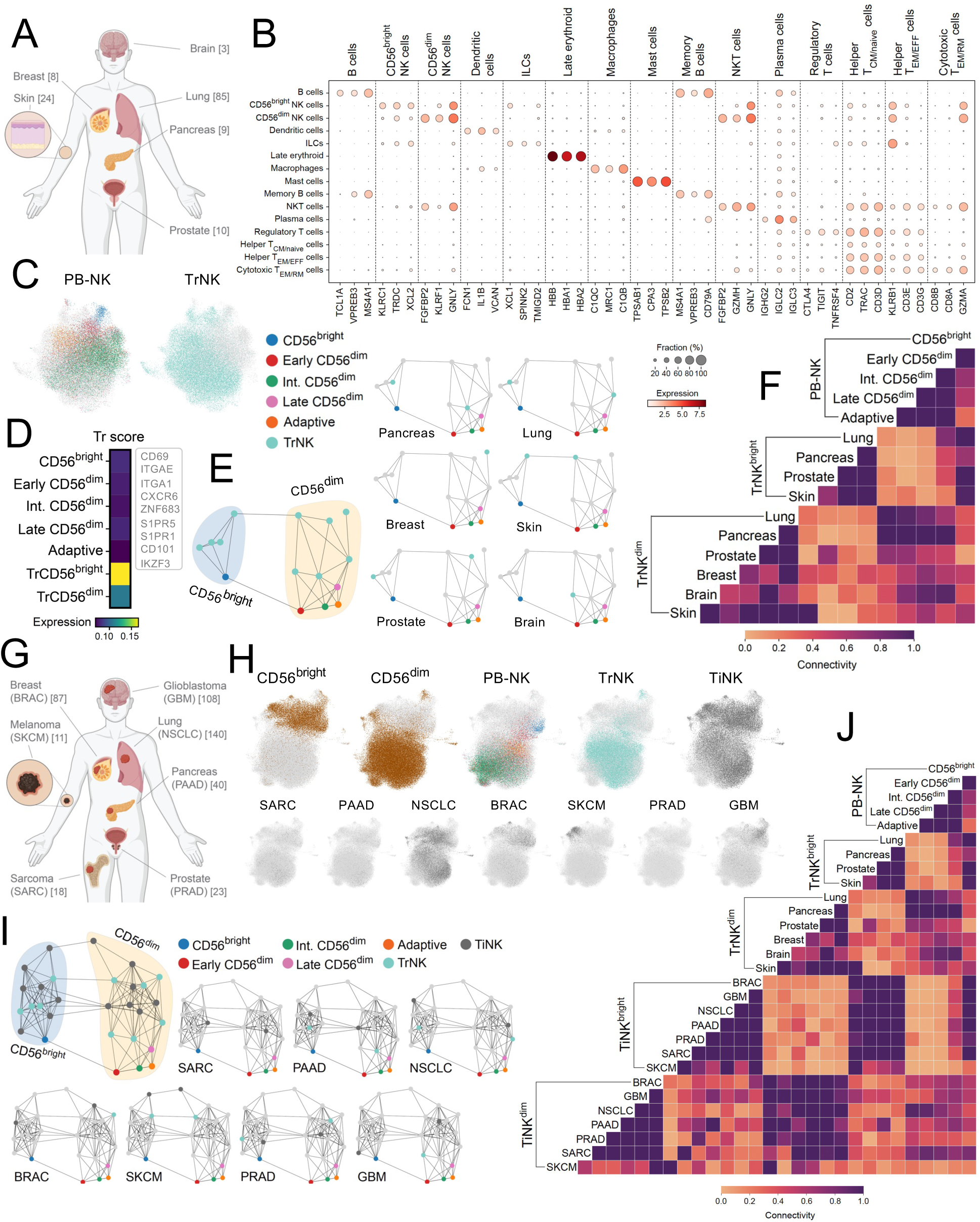
Pan-cancer atlas of healthy tissue resident and solid tumor-infiltrating NK cells. (**A**) Graphical overview of healthy tissue datasets included in the analysis, with the number of donors denoted in brackets. (**B**) Dotplot showing selected signature genes, and their expression in healthy lung, used to annotate major immune subsets in the compiled dataset. (**C**) UMAP representation showing integration of subset annotated peripheral blood-derived (PB-NK) and tissue-derived NK cells (TrNK). (**D**) Scoring of tissue residency signature in PB-NK cell subsets and CD56^bright^ and CD56^dim^ annotated TrNK subsets. (**E-F**) PAGA graph (**E**) and connectivity heatmap (**F**) showing connectivity of PB-NK and TrNK subsets across all tissues, with individual tissues highlighted (**E**). (**G**) Graphical overview of solid tumor datasets included in the analysis, with the number of donors denoted in brackets. (**H**) UMAP representation showing integration of subset annotated PB-NK, TrNK and tumor-infiltrating NK cells (TiNK) as pan-cancer atlas and stratified by solid-tumor type. (**I-J**) PAGA graph (**I**) and connectivity heatmap (**J**) showing connectivity of PB-NK, TrNK and TiNK subsets across all tissues/tumor types, with individual tissue/tumor types highlighted.

Next, scRNA-seq datasets from seven solid tumors (breast cancer (BRAC), Glioblastoma (GBM), Lung (NSCLC), Melanoma (SKCM), Pancreas (PAAD), Prostate (PRAD) and Osteosarcoma (SARC)) from a total of 427 patients were annotated and integrated for each tumor type using scANVI^19^ and CellTypist^39^ (**Figure 3G, Supplemental Figure 3F-L, Supplemental Table 3**). The CD56^bright^ and CD56^dim^ annotated tumor-infiltrating NK (TiNK) cells were mapped onto the reference map (PB-NK, TrNK) using transfer learning (scArches^40^) to generate the final model (M3) (**Figure 3H**). TiNK cells also clustered based on their differentiation stage together with the corresponding PB-NK and TrNK subsets (**Figure 3I**). SKCM-derived CD56^dim^ NK cells exhibited the lowest connectivity score when compared to all other populations (**Figure 3J**). GBM-derived TiCD56^bright^ NK cells and SKCM-derived TiCD56^dim^ scored highest for tissue residency within their respective subsets (**Supplemental Figure 3M**). Transfer learning facilitated incorporation of TiNK cells onto our healthy reference map of PB and TrNK cells, allowing for downstream systematic interrogation of cellular states within these solid-tumor infiltrating NK cells.

### Altered NK cell subset frequencies within healthy tissue and solid tumors

The tumor microenvironment (TME) is shaped by its cellular composition, particular by the infiltrating immune cells, which in turn can be modulated by their surroundings. A pan-cancer comparison of the healthy tissue and tumor annotated immune subtypes (**Figure 3B**), identified an increased proportion of plasma cells and a decreased proportion of CD56^dim^ NK cells, dendritic cells, NKT cells, helper TEM/EFF, cytotoxic TEM/EMRA and cytotoxic TRM cells in the tumor datasets (**Figure 4A-B**). The fraction of CD56^bright^ NK cells out of total immune cells was enriched in BRAC, while CD56^dim^ NK cells were enriched in SKCM, but decreased in NSCLC and BRAC (**Figure 4B, Supplemental Figure 4A**). SKCM uniquely exhibited a tendency for increased proportions of both NK cell subsets (**Figure 4B**), in line with an overall increased frequency of immune cells, including NK cells (**Figure 4C-D**). Utilizing our subset-trained model (M1) to annotate CellTypist defined NK cells into the five NK cell subsets we observed increased proportions of CD56^bright^ cells across numerous tumor types (**Figure 4E**). Within the CD56^dim^ compartment, a skewing towards more mature NK cells (late CD56^dim^) in tissues and tumors compared to blood (intermediate CD56^dim^) was detected, with adaptive NK cells notably absent in Tr/TiNK cells (**Figure 4E**). Solid tumor-infiltrating NK cells were enriched for a CD56^bright^ transcriptional phenotype while the CD56^dim^ compartment in both healthy tissue and solid tumors was skewed towards increased maturity (late CD56^dim^).

**Figure 4.**
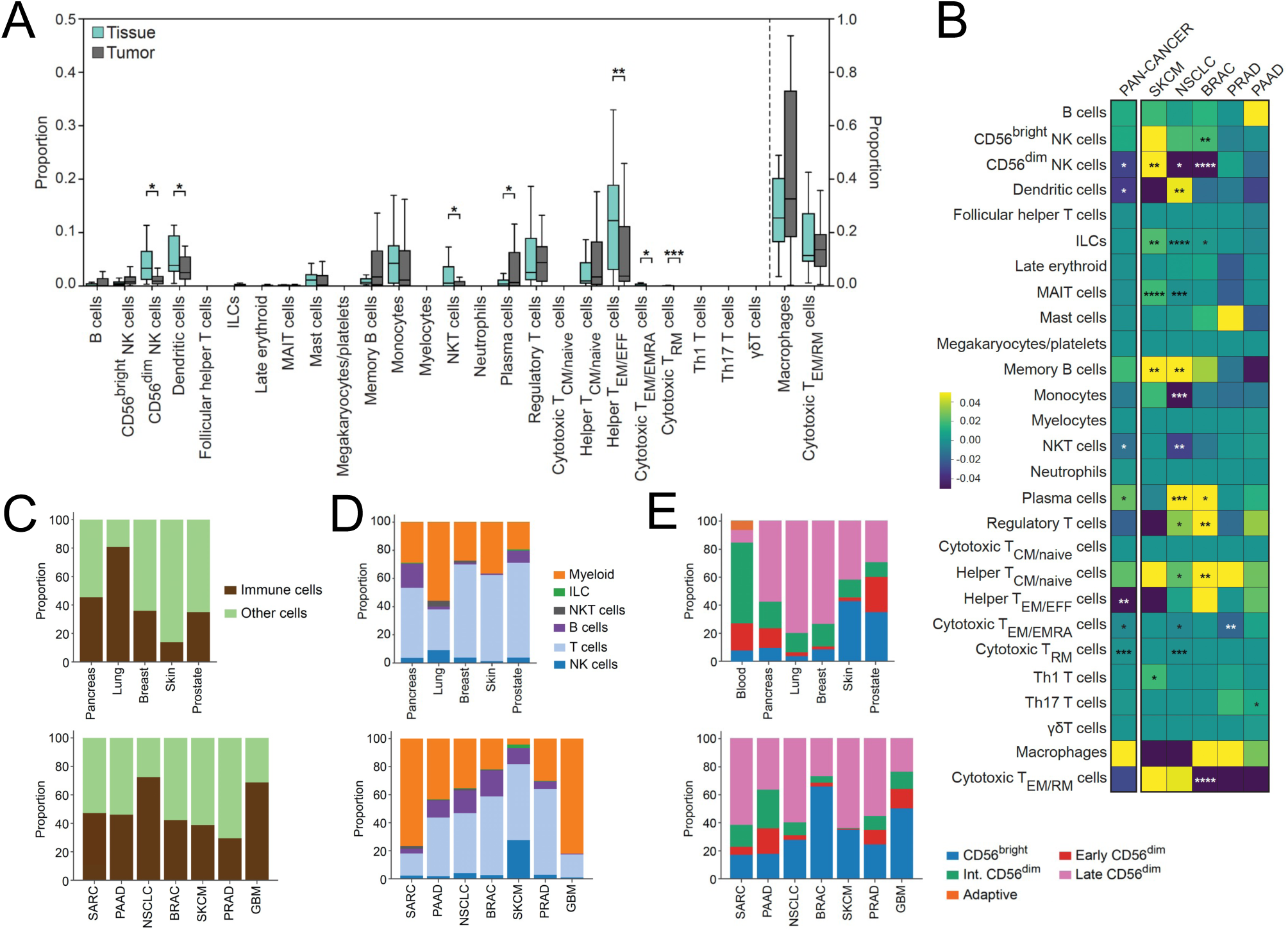
Cellular composition of pan-cancer cell atlas and subset distribution of tumor- infiltrating NK cells. (**A**) Distribution of major immune subsets across all tissue and tumor types. (**B**) Heatmap depicting changes in immune subset proportion in tumor samples compared to healthy tissue samples at the pan-cancer level and within individual tumor types. (**C**) Proportion of immune cells out of total cells within healthy tissue samples and tumor samples. (**D**) Proportions of major immune subsets within healthy tissue and tumor samples. (**E**) Predicted subset annotations of CellTypist identified NK cells in healthy tissue and tumor samples compared to annotated PB-NK cells. Boxplots (center line – median, box limits – upper/lower quartiles, whiskers – distribution). Data were analyzed using two-sample t-test with Bonferroni correction; * p < 0.05, ** p < 0.01, *** p < 0.001, **** p < 0.0001.

### Six distinct functional states of NK cells in peripheral blood, tissues, and tumors

Tumor microenvironments of solid tumors are hostile and often immunosuppressive environments for immune cells to infiltrate.^41^ Understanding how the TME can modulate NK cells at the transcriptional level can provide important insights into understanding the tumor-mediated immunosuppressive mechanisms and how to overcome them.

We implemented an unbiased approach (Milo^42^) to ascertain cellular states in our pan-cancer NK cell atlas by identifying individual neighborhoods (∼6000) without pre-clustering based on cellular origin. Annotating individual neighborhoods as subset specific (>70% of cells in neighborhood) identified TiCD56^bright^ NK cells as having the most frequent, but also most unique (differentially abundant) specific neighborhoods (**Supplemental Figure 5A**). Notably, the majority of neighborhoods were annotated as ‘mixed’, highlighting transcriptional similarities among NK cells found in peripheral blood, tissues and tumors (**Supplemental Figure 5A**). The approximately 6000 neighborhoods were grouped into six distinctive neighborhood groups and we tested for differential abundance of neighborhoods between TiNK cells and Ref-NK cells (**Figure 5A, Supplemental Figure 5B**). Neighborhood groups 1 and 2 consisted of neighborhoods significantly enriched for TiNK cells and group 6 included neighborhoods enriched for Ref-NK cells (**Figure 5B, Supplemental Figure 5B**).

**Figure 5.**
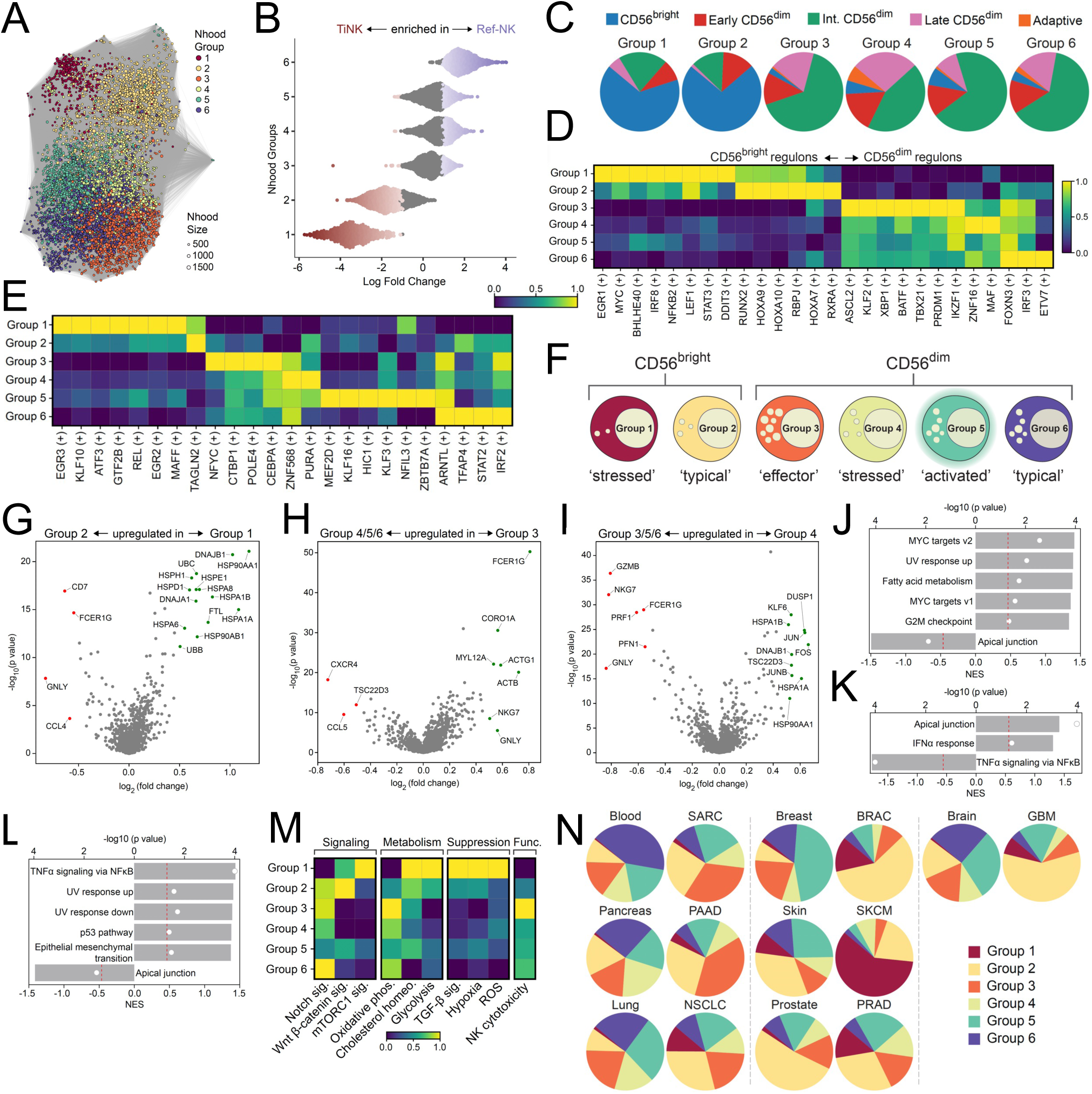
Distinct cellular states of NK cells identified in pan-cancer atlas. (**A**) UMAP depicting neighborhood groups identified by Milo. (**B**) Beaswarm plot depicting differential abundance of neighborhoods (TiNK vs Ref-NK enriched). Colored neighborhoods are differentially abundant at FDR 0.1. (**C**) Pie charts showing distribution of NK subsets across neighborhood groups annotated using our annotation Model (Figure 1). (**D**) Expression of dominant transcription factor (TF) regulons of NK cell differentiation across NK cell states (neighborhood groups). (**E**) Expression of TF regulons uniquely expressed across cellular states. (**F**) Graphical representation of cellular states. (**G-L**) Volcano plots depicting differentially expressed genes (DEGs) and corresponding gene set enrichment analysis (GSEA) between Group 1 vs. Group 2 (**G, J**), Group 3 vs. Group 4/5/6 (**H, K**) and Group 4 vs. Group 3/5/6 (**I, L**) cellular states. (**M**) Scoring of pathway gene signatures in NK cells states. (**N**) Pie charts depicting distribution of NK cell states in blood, tissues and tumors. Volcano plots: log fold change cutoff at 0.5, p < 0.05. GSEA plots: p value cutoff 0.5 (red line).

Next, we visualized the distribution of NK cell subsets within each group using our annotation model (M1). Group 1 and 2 were enriched for, but not exclusive to CD56^bright^ cells, while groups 3-6 were dominated by CD56^dim^ NK cell subsets (**Figure 5C**). The dominant TF regulons of PB-NK cell differentiation previously identified (**Figure 2F**), confirmed Group 1 and 2 as two CD56^bright^ states and group 3-6 as four CD56^dim^ NK cells states (**Figure 5D**).

Cell-state specific GRN, DEG, GSEA, and signature scoring informed our annotation of the states as ‘stressed’ CD56^bright^ (Group 1), ‘typical’ CD56^bright^ (Group 2), ‘effector’ CD56^dim^ (Group 3), ‘stressed’ CD56^dim^ (Group 4), ‘activated’ CD56^dim^ (Group 5) and ‘typical’ CD56^dim^ (Group 6) (**Figure 5E-M**, **Supplemental Figure 5C-F**). Comparing the ‘stressed’ to the ‘typical’ CD56^bright^ state identified increased expression of the cellular stress response ATF3 regulon, the hypoxia-induced MAFF regulon, and numerous heat shock proteins (**Figure 5E**, **G, J**). The ‘stressed’ CD56^bright^ cell state scored highly for immunosuppressive pathways (TGF-β signaling, hypoxia, ROS) and exhibited increased metabolic activation (glycolysis, cholesterol homeostasis, fatty acid metabolism), proliferation (G2M checkpoint) and activation of the MYC/mTORC1 axis (**Figure 5G, J, M**). Furthermore, a significant decrease in the apical junction hallmark (indicative of lower polarization and conjugate formation) and a low NK cytotoxicity score was suggestive of reduced functionality in this ‘stressed’ CD56^bright^ cellular state, which was uniquely enriched across all 7 tumor types (**Figure 5J, M-N**). In line with increased infiltration of CD56^bright^ cells in the TME, the ‘typical’ CD56^bright^ cellular state was also enriched in 5 of 7 tumor types compared to healthy tissue (**Figure 5N**).

Of the CD56^dim^ state, the ‘effector’ state was most frequently enriched across tumor types (SARC, PAAD, PRAD), characterized by an enrichment for apical junction, actin and cytoskeleton-related associated genes (**Figure 5H, K, N**). This state, phenotypically enriched for intermediate and late CD56^dim^ NK cell subsets, scored highly for NK cytotoxicity and oxidative phosphorylation, and importantly, lowly for immune suppression (**Figure 5C, M**). The ‘stressed’ CD56^dim^ state, characterized by downregulated apical junction related genes and effector molecules (GZMB, PRF1, GNLY) and upregulated heat shock proteins, was more prominent in healthy tissues and only enriched for in PRAD (**Figure 5I, L-N**). The ‘activated’ CD56^dim^ state was distinguished by increased hypoxia, proliferation and NFκB activation (**Supplemental Figure 5C, E**, **Figure 5M**) while the PB-enriched ‘typical’ CD56^dim^ state exhibited highest expression of NK-associated genes (PRF1, GZMB, CST7, FCGR3A, NKG7, FGFBP2) (**Supplemental Figure 5D, F**, **Figure 5N**). Notably, while we observed enrichment of individual cellular states in the TME, including the two CD56^bright^ and the ‘effector’ CD56^dim^ states, all states were represented in healthy blood and tissue samples, albeit at different frequencies.

### Decreased TME-specific incoming signaling in the ‘effector’ CD56^dim^ NK state associated with improved survival

The clinical benefit of NK cell infiltration in solid tumors has previously been assessed through a general NK cell signature score^43, 44^. Having identified six functional states of NK cells in blood, tissue and solid tumors, we proceeded to test clinical relevance of these cellular states by using BayesPRISM^45^ to deconvoluted TCGA survival data^46, 47^. The combination of high ‘effector’ CD56^dim^ and low ‘stressed’ CD56^bright^ cell signatures correlated with increased improved survival in SARC and SKCM patients (**Figure 6A**). To elucidate any TME-based influence on these outcome-associated functional states, we employed CellChat^48^ to infer intercellular communication, focusing on commonly enriched signaling pathways in SARC and SKCM. Increased outgoing signaling (MHC-I, CD99, ITGB2, ICAM, PARs) was noted in group 3 NK cells, while group 1 NK cells were enriched for incoming signaling (MHC-I, MIF, ADGRE5, FN1, GALECTIN, COLLAGEN) (**Figure 6B, Supplemental Figure 6A**). Increased expression of CD44, and to a lesser degree CXCR4, upon which numerous signals from fibroblasts, CAFs, endothelial cells and osteoblasts/clasts converged (COLLAGEN, MIF GALECTIN; FN1), facilitated the augmented incoming signaling in group 1 (**Figure 5C, E**). Notably, fibroblasts, CAFs, endothelial cells and osteoblasts/clasts also exhibited the strongest outgoing interaction strength of all cell types in SARC (**Supplemental Figure 6A**). Furthermore, group 1 NK cells preferentially received inhibitory input via the MHC-I (HLA-E/KLRC1) and ADGRE5 (ADGRE5/CD55) pathways, while group 3 NK cells exhibited increased ITGB2 and ICAM2 expression, facilitating binding to other NK cell states and macrophages (**Figure 6D-E**). Hence, group 3 NK cells preferentially communicated with other tumor-infiltrating immune cells while group 1 NK cells were more receptive to TME-induced immunosuppressive signals via upregulated CD44.

**Figure 6.**
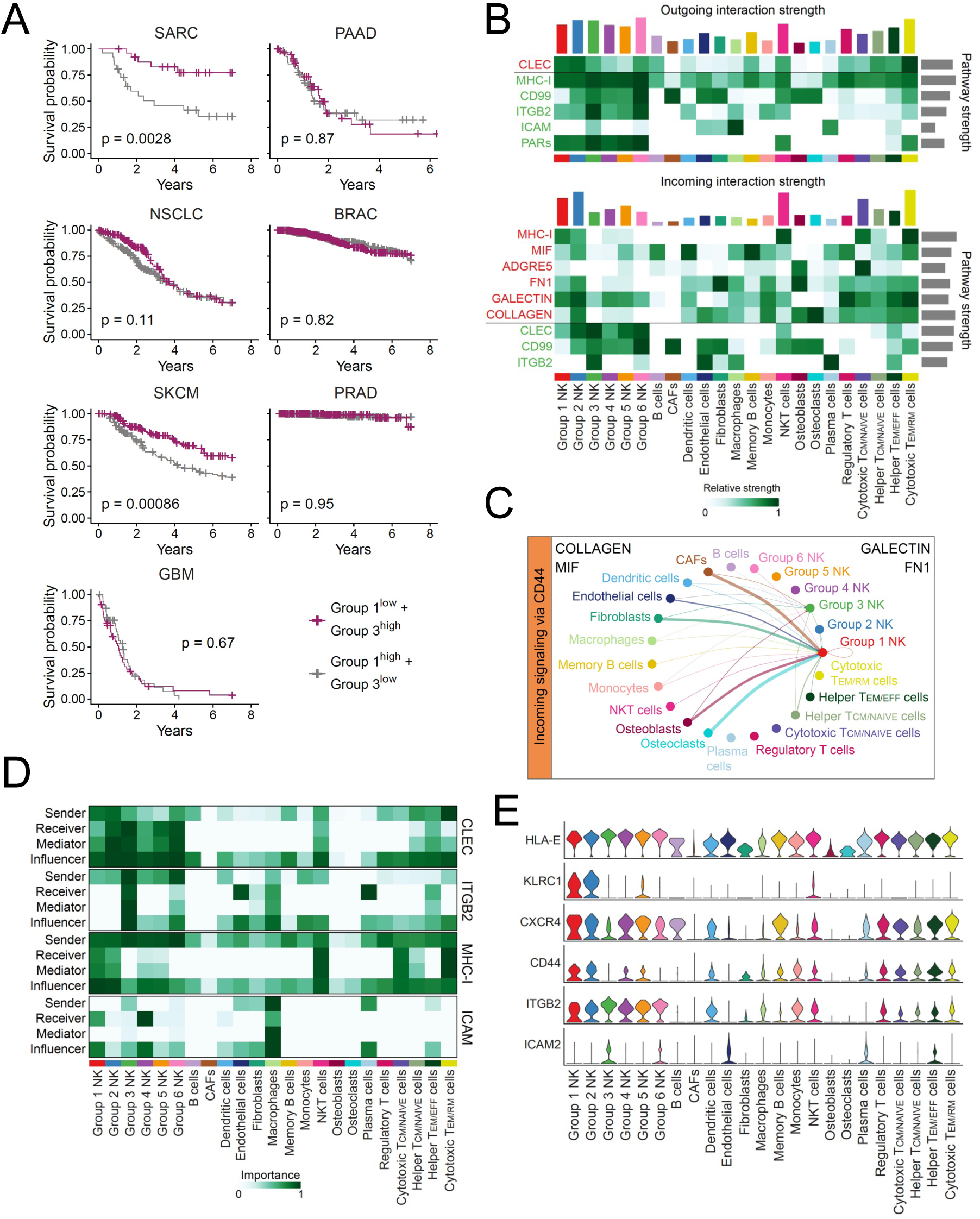
Intercellular communication of distinct cellular states associated with patient outcome. (**A**) Kaplan-Meier survival curves showing association of high/low Group 1/3 gene signatures with patient outcome across tumor types. (**B**) Selected predicted outgoing (top) and incoming (bottom) signaling pathways involving TiNK cells in SARC as identified by CellChat. Pathways in red are enriched for in Group 1 NK cells and pathways in green are enriched for in Group 3 NK cells. (**C**) Circle plot depicting predicted incoming signaling via CD44 expression on Group 1 and Group 3 TiNK cells (SARC). (**D**) Heatmap depicting importance and interaction role of individual cell populations in CLEC, ITGB2, MHC-1 and ICAM signaling pathways in SARC based on network centrality analysis in. (**E**) Violin plots showing expression of receptors and ligands of communication pathways involving TiNK cells in SARC. MHC-I (HLA-E – KLRC1), ITGB2 (ICAM2 – ITGB2), COLLAGEN/GALECTIN/FN1 (CD44), MIF (CD44+CXCR4).

A higher ratio of ‘effector’ CD56^dim^ to ‘stressed’ CD56^bright^ NK state signatures was predictive of improved survival in SARC and SKCM. Inferred increased inhibitory signaling and augmented susceptibility to TME-induced suppression likely contributes to the ‘stressed’ CD56^bright^ states unfavorable prognosis.

## Discussion

Here we report a compact description of the transcriptional diversification encompassing human NK cell differentiation at the single cell level. By enriching for less frequent, but phenotypically well-defined functionally distinct NK cell subsets, we could first train a model to correctly annotate five transcriptional subsets from bulk NK cell populations. By applying probabilistic models implemented in scVI-tools, we created a transcriptional reference map of human blood and tissue resident NK (TrNK) cells from normal tissues including blood, pancreas, lung, breast, skin, prostate and brain. Transfer learning using scArches facilitated integration of query datasets comprising a total of 2,176,214 transcriptomes from 427 patients spanning seven solid tumor types. By extracting, annotating, and mapping the tumor-infiltrating NK (TiNK) cells onto our healthy reference map, we could systematically interrogate TME-induced perturbations of gene regulatory networks and functional states of TiNK cells (**Supplemental Figure 7**). Our pan-cancer atlas revealed six functionally distinct NK cell states with varying abundance across blood, tissues and tumor types. Two states commonly enriched for across tumor types included a dysfunctional CD56^bright^ cellular state susceptible to TME-induced immunosuppression and a cytotoxic TME- resistant CD56^dim^ state, the ratio of which was predictive of patient outcome.

The view that NK cells, like T cells and other immune cells, undergo a continuous process of NK cell differentiation is relatively recent and was originally based on phenotypic and functional classification of discrete subsets^7, 49^. There is abundant evidence suggesting that the CD56^bright^ NK cell subset is the most naïve, giving rise to the more differentiated CD56^dim^ NK cells which can further differentiate towards terminal stages, a process accelerated by CMV infection^8, 50, 51^. Instead of forcing individual NK cells into arbitrary clusters representing a snapshot of a given time point of differentiation, we clustered TFs and their target genes into five distinct gene expression trends as a function pseudotime, reflecting continuous differentiation. By retaining fate-specific expression profiles, conventional versus adaptive fate in donors with CMV-induced clonal NK cell expansions, we could observe clear divergence of regulon expression (eg, BATF, MAF) during terminal differentiation. BATF belongs to the AP-1 TF family which have been identified as potential drivers in shaping adaptive NK cell chromatin accessibility and thus dictating the unique functional features of this subset, including enhanced IFNγ response to receptor stimulation^15^. Establishing dominant regulons defining NK cell differentiation in peripheral blood provided a vital reference for downstream interrogation of both tissue resident and solid tumor-infiltrating NK cells.

Utilizing CellTypist, we harmonized annotations of individual cell subtypes across multiple datasets from six different healthy tissues, extracting and integrating CD56^bright^ and CD56^dim^ NK cells using scVI^19^ to expand our transcriptional reference map. Importantly, tissue, as well as tumor-annotated NK cells did not express human ILC signature genes, instead expressing both EOMES and TBX21. Tissue residency genes (e.g., CD69, ITGAE, ITGA1, CXCR6, ZNF683, IKZF3) were more highly expressed in tissue-derived NK cells, particularly in CD56^bright^ NK cells. Notably, we could not identify a CD56^bright^ population in both healthy brain and breast datasets. This could be attributed to the absence of CX3CR1^52^ expression in CD56^bright^ NK cells, an important receptor for NK cell migration to the brain, or could be an artefact due to higher blood contamination (lower CD56^bright^ frequency) in this dataset in line with a lower tissue residency score.

The presence and abundance of NK cells that reside in the tumor bed varies across tumor types, treatments and between patients and appears to be associated with the chemokine profiles in the different tissues/tumor microenvironments^53, 54, 55, 56^. Immune and NK cell subset composition greatly varied among tissue and tumor type, with the highest and lowest frequency of CD56^bright^ NK cells being found in skin and lung respectively. Consistently across tissue and tumor type, a clear maturation of the CD56^dim^ subset was noted, with late CD56^dim^ NK cells making up the largest fraction. Notably, no Tr nor TiNK cells were annotated as adaptive using our subset annotation model, confirmed by Tang et al.^16^ but contrary to previous reports describing adaptive-like NK cells with a tissue-residency phenotype in the lung^57^. Transcriptional differences between previously described tissue-resident adaptive NK cells and our PB-derived gene signature trained annotation model could explain these discrepancies.

In agreement with previous studies^54, 58^, we observed a predominance of CD56^bright^ NK cells in tumors compared to the corresponding normal tissue. Tumor-resident NK cells are likely a mixed population including naturally residing TrNK cells and TiNK cells. Compositional differences between normal and tumor tissues suggests some degree of active recruitment, particularly in SKCM where NK cell frequencies starkly increased, albeit expansion from tissue resident pools cannot be excluded. Migration into the TME is regulated by a broad family of integrins, selectins and chemokine receptors that are differentially expressed during NK cell differentiation. CXCR3, primarily expressed CD56^bright^ NK cells, has been implicated in homing to several solid tumors based on CXCL10 gradients^59^, and thus may contribute to the predominance of this subset in tumors. CCL2, CCL3, CCL5, CXCL8, CXCL9, CXCL10, and CXCL12, have similarly been implicated in mediating predominantly CD56^bright^ NK cell trafficking into solid tumors based on chemokine receptor expression^56^. We observed heightened CXCR4 expression in CD56^bright^ Tr and TiNK cells, and a modest upregulation of CX3CR1 on CD56^dim^ Tr and TiNK cells, with levels varying across tissue/tumor type. Previous reports^60, 61^ have demonstrated CD44-induced CXCR4 upregulation resulting in increased migration and invasiveness of malignant cells. Notably, CD44 was highly expressed on the tumor-enriched ‘stressed’ CD56^bright^ state, possibly sensitizing this population to TME-mediated immunosuppression from CAFs, fibroblasts, endothelial and tumor cells, as noted by high scores for TGFβ signaling, hypoxia and ROS. Conversely, the ‘effector’ CD56^dim^ state associating with improved patient outcome, lacked CD44 expression and uniquely expressed ICAM2 and high ITGB2 levels. Notably, this state exhibited high expression of the KLF2, PRDM1, BATF, TBX21 and IKZF1 regulons, indicative of high effector function, regulation of homeostatic proliferation and survival, but also cell migration and tissue residency. Unique TiNK specific regulons in this state consisted of NFYC, CTBP1, POLE4 and CEBPA, which are involved in DNA repair, monitoring of proliferation, regulating MHC expression and maintaining structural homeostasis in the Golgi complex^62, 63, 64, 65^. Conversely, TiNK specific regulons in the ‘stressed’ CD56^bright^ state included hypoxia induced MAFF, cellular stress response regulon ATF3 and EGR2/3^66^ which induce negative regulators in response to activation. Contrary to Tang et al.^16^, increased gene signature scoring of the tumor enriched ‘stressed’ CD56^bright^ state did not consistently associate with reduced survival across tumor types. Instead, we observed increased survival in patients exhibiting a high ‘effector’ CD56^dim^ state which was further augmented with a low signature for the ‘stressed’ CD56^bright^ state. Of the four CD56^dim^ states, the ‘effector’ CD56^dim^ state was most commonly enriched across tumor types, painting a promising picture for the role of solid-tumor infiltrating NK cells.

This resource provides a transcriptional reference map of human NK cells across healthy blood and tissues with harmonized annotations of transcriptional NK cell subsets. Uncovering the dominant gene regulatory circuits during NK cell differentiation enabled identification of TME-induced perturbations in solid tumor-infiltrating NK cells across tumor type. We identified functionally distinct NK cell states across healthy and malignant tissues, including tumor enriched states predictive of patient outcome. Modelling of the intercellular communication pathways of outcome-predicting NK cell states with the surrounding TME identified potential pathways of TME-induced NK cell suppression. Thus, our analysis has the potential to design more potent NK cell therapy products able to resist suppressive factors operating within the TME of solid tumors. Ultimately, this resource can be extended endlessly through transfer learning to interrogate new datasets from experimental perturbations or different tumor types.

## Acknowledgements

Large parts of the analyses were run using the Machine learning infrastructure (ML Nodes), University Centre for Information Technology, University of Oslo, Norway. This publication is part of the Human Cell Atlas –www.humancellatlas.org/publications/

## Data availability

The gene expression data generated for this paper is available at NCBI GEO with accession number GSE245690 and raw sequencing data is available at EGA with accession number EGAS50000000014. The details about the publicly available data included in the analysis are available in **Supplemental tables S1, S2** and **S3**. Processed data and models have also been made available on Zenodo (https://doi.org/10.5281/zenodo.8434224) and as an online resource at http://nk-scrna.malmberglab.com/.

## Code availability

The code generated for our analysis is available on GitHub at http://github.com/hernet/transcriptional-map-nk

## Authorship and conflict-of-interest statements

J.G, A.H and A.P. performed the single-cell RNA sequencing experiments. H. N., A.P. and T. C. performed the bioinformatic analysis. E.S., O.D., S.A.T., A.H. and K-J.M. provided scientific input. A.P. H.N, and K-J.M. wrote the manuscript. J.G. is an employee at Fate Therapeutics. K-J.M. is a consultant at Fate Therapeutics and Vycellix and has research support from Fate Therapeutics, Oncopeptides for studies unrelated to this work. S.A.T. is a co-founder and board member of and holds equity in Transition Bio. Figures were partly generated using Biorender software.

Survival analysis was performed using the Cox proportional hazards model, p values were computed using the log-rank test.

## Methods

### Cell processing

Peripheral mononuclear cells (PBMC) were isolated using density gradient centrifugation from anonymized healthy blood donors (Oslo University Hospital; Karolinska University Hospital) with informed consent. The study was approved by the regional ethics committee in Norway (2018/2482) and Sweden (2016/1415-32, 2020-05289). Donor-derived PBMCs were screened for KIR education and adaptive status using flow cytometry. NK cells were purified using an AutoMACS (DepleteS program, Miltenyi Biotec) and prior to overnight resting in complete RPMI (10% Fetal calf serum, 2mM L-glutamine) at 37°C/5% CO2.

### Flow cytometry screening

PBMC were stained for surface antigens and viability in a 96 V-bottom plate, followed by fixation/permeabilization and intracellular staining at 4°C. The following antibodies were used in the screening panel: CD3-V500 (UCHT1), CD14-V500 (MφP9), CD19-V500 (HIB19), Granzyme B-AF700 (GB11) from Beckton Dickinson; CD57-FITC (HNK-1), CD38-BV650 (HB-7), KIR3DL1-BV421 (DX9) from BioLegend; KIR2DL1-APC-Cy7 (REA284), CD158a,h-PE-Cy7 (11PB6), from Miltenyi Biotec; CD158b1/b2,j-PE-Cy5.5 (GL183), NKG2A-APC (Z199), CD56-ECD (N901) from Beckman Coulter. LIVE/DEAD Fixable Aqua Dead Stain kit for 405 nM excitation (Life Technologies) was used to determine viability. Samples were acquired on an LSR-Fortessa equipped with a blue, red and violet laser and analyzed in FlowJo version 9 (TreeStar, Inc.).

### FACS sorting

Cells were harvested and surface stained with the following antibodies: CD57-FITC (HNK-1) from BioLegend; KIR3DL1S1-APC (Z27.3.7), CD56-ECD (N901), CD158b1/b2,j-PE-Cy5.5 (GL183), from Beckman Coulter; KIR2DL1-APC-Cy7 (REA284), NKG2C-PE (REA205), NKG2A-PE Vio770 (REA110) from Miltenyi Biotec. 12,000 cells were directly sorted into Eppendorf tubes at 4°C for each sample using a FACSAriaII (Beckton Dickinson). Sorting strategies for single-cell RNA sequencing for the donor with an adaptive NK cell expansion and without are depicted in **Supplemental Figure 1C and 1D** respectively.

### Single-cell RNA sequencing

Following sorting, cells were kept on ice during the washing (PBS + 0.05% BSA) and counting step. 10,000 cells were resuspended in 35 μL (PBS + 0.05% BSA) and immediately processed at the Genomics Core Facility (Oslo University Hospital) using the Chromium Single Cell 3’ Library & Gel Bead Kit v2 (Chromium Controller System, 10X Genomics). The recommended 10x Genomics protocol was used to generate the sequencing libraries, which was performed on a NextSeq500 (Illumina) with 5∼ % PhiX as spike-inn. Sequencing raw data were converted into fastq files by running the Illuminàs bcl2fastq v2.

### ScRNAseq data collection and processing

Previously published scRNA-seq data were collected mostly in the form of count matrices already aligned to GRCh38, the rest was collected as fastq files. For the datasets where we collected fastq files, the data was aligned to GRCh38 using Cell Ranger (10x Genomics Cell Ranger 7.0.0).

### Quality control and normalization of scRNA-seq data

Data cleaning steps were first carried out whereby cells not expressing a minimum of 1000 molecules and genes expressed by less than 10 cells were filtered out. Doublets were removed using the SOLO algorithm^1^. The data was normalized using log transformation for some of the downstream analysis as well as for visualization of gene expression like dot plots. Quality control, transformation and most of the visualization of the gene expression data was performed using Scanpy^2^. For analysis using scVI and scANVI the raw count data was used.

### Integration of scRNA-seq data

The probabilistic models scVI and scANVI as implemented in scvi-tools^3^ were used for integration of scRNA-seq data. These methods have been shown to perform well for integration of scRNA-seq data, especially when dealing with complex batch effects and integrating atlas- level data^4^. For cell type and subset annotations and prediction scANVI was used to capture annotation of single-cell profiles. For the analysis of PB-NK subsets the sorted subsets provided labels for training the scANVI model. The subset prediction provided by the model was tested on a held out set of cells (15%) from the sorted subset data giving us a confusion matrix summarizing the performance of the prediction.

### Dimensionality reduction, clustering and visualization of scRNA-seq data

We computed the UMAP embeddings for visualization using the embedding learned from scVI and scANVI. Unsupervised clustering was also carried out using this learnt embedding using the Leiden algorithm as implemented in Scanpy. PAGA^5^ was used to quantify the connectivity of different groups of cells and thereby providing a representation of the data as a simpler graph. The various plots were mostly generated using the plotting functions in Scanpy.

### Cell type annotations and harmonization

For many of the publicly available datasets cell type annotations were readily available and used as seed labels when training the scANVI model for that particular tissue/tumor type. The scANVI model allowed us to harmonize annotations which was needed for analysis across datasets. Celltypist^6^ was also used for annotations, specifically for the immune cell compartment in the various tissue/tumor types. The CD16- and CD16+ NK cells identified by Celltypist were annotated as CD56^bright^ and CD56^dim^ respectively. Where CITE-seq data was available the surface expression of key markers also helped validate the cell type annotations. For the identified NK cells the cells were also scored using NK1/NK2 (CD56^bright^/CD56^dim^) signatures to validate the annotation of CD56^bright^ and CD56^dim^ NK cells. We also performed our own unsupervised Leiden clustering which identified two dominating clusters corresponding to CD56^bright^ and CD56^dim^ NK cells.

### Calculation of signature scores

Signature scores were computed using AUCell^7^ allowing for exploration of the relative expression of the signatures of interest in the data sets. Various gene sets were taken from the MSigDB Hallmark gene set collection^8^.

### Pseudotime and RNA velocity analysis

Pseudotime was computed using Palantir^9^ which captures the continuous nature of differentiation and cell fate which allowed us to explore two terminal states and the gene expression changes seen along these trajectories. For this analysis the starting cell was defined as the cell that was the least CD56^dim^ (the lowest score for the NK1 signature). Generalized- additive models (GAMs) fitted on cells ordered by pseudotime were used to calculate gene trends, where the contribution of cells was weighted by their probability to end up in the given terminal state as calculated by Palantir. The gene trends indicate how gene expression levels develop over the differentiation timeline. These trends were clustered using the Leiden clustering algorithm to give us five clusters of gene trends. RNA velocity^10^ was also used in order to take advantage splicing kinetics to identify directed dynamic information. We used velocyto^10^ and scVelo^11^ for this analysis, specifically the dynamic model implemented in the scVelo toolkit. The RNA velocity analysis was run on the two donors where sorted subsets where sequenced separately, as well as on the integrated data from 12 blood donors.

### Gene regulatory network analysis

SCENIC^7^ was used to infer transcription factors and gene regulatory networks from the scRNA-seq data. The SCENIC workflow^12^ was followed and the pySCENIC implementation was used. TF-gene associations were inferred by GRNBoost^13^ and motif-to-TF associations were downloaded from the Aerts’s lab website and used for pruning the inferred associations. The inferred regulatory networks were also further pruned by removing lowly expressed TFs based on the bulk RNA-seq data. AUCell was used to compute the activity of the final regulons. The regulon activity was visualized using matrix plots as implemented in Scanpy to look at the activity across different groups of cells.

### Bulk RNA sequencing for TF and target validation

For validation of the TF and targets we checked their expression in bulk RNASeq data from four sorted NK cell populations (CD56^bright^, NKG2A^-^KIR^-^CD56^dim^, NKG2A^-^KIR^+^CD56^dim^, and NKG2A^-^KIR^+^NKG2C^+^CD56^dim^). Sequencing was performed using single-cell tagged reverse transcription (STRT)^14^.

### Reference mapping

The TiNK cells were added after the model for a healthy NK cell reference was trained. scArches^15^ as implemented in scvi-tools^3^ was used to map this new data onto the established reference.

### Cell-cell communication inference using CellChat

To infer the communication between the various cell types in the tumor data sets we used CellChat^16^. Based on gene expression of receptors and ligands in the data and a curated database of pathways, CellChat computes the communication probability between various receptor-ligand pairs. CellChat also provided ways to aggregated this information and for us to visualize the inferred cell-cell communication networks. CellChat was computed separately for each of the tumor types included in the analysis.

### Differential gene expression analysis

In order to perform differential gene expression analysis we used pseudobulk as this has shown good results when analyzing scRNA-seq data in various studies^17^. This allowed us to aggregate up counts for each sample and consider the samples instead of the cells as replicates. We then used edgeR^18^ on the pseudobulk data. We could then identify differentially expressed genes between healthy reference NK cells and TiNK cells within and across subsets.

### Differential abundance analysis using Milo

We used Milo^19^ to assign cells to neighborhoods on the KNN graph. The differential abundance of these neighborhoods between the healthy reference and the TiNK cells were then computed. The neighborhoods were grouped into six groups using the *groupNhoods* function in Milo. These groups were considered as different NK cell states and further characterized using the functions in Milo for identification of differentially expressed genes. The single cells were also annotated using these groups for downstream analysis.

### Gene set enrichment analysis

Gene set enrichment analysis was performed using the GSEA software^20^ and the MSigDB collection of gene sets. Genes were first ordered based on the differential expression analysis either based on the pseudobulk approach or based on the Milo analysis.

### Clinical and bulk RNA-seq data from TCGA and TARGET

Bulk RNA-seq data and clinical data was downloaded from TCGA and TARGET using TCGAbiolinks^21^ and curated survival data was downloaded from Xena^22^.

### Deconvolution of bulk RNA-seq

Deconvolution of the bulk RNA-seq data was performed for each of the tumor types using BayesPrism^23^. BayesPrism has been shown to work well for deconvolution of data from tumors and especially well in dealing with high cell type granularity^24^. The annotated reference datasets for each of the tumor types were used as prior information in the deconvolution. BayesPrism then computed both an expression matrix for each cell type as well as the cell type fraction for each sample.

### Survival analysis

The NK expression matrix inferred by BayesPrism for the various tumor types were used to score the signature genes for each of the identified NK cell states. The patients were then assigned as high and low for a group/state based on belonging to the highest or lowest half in terms of expression of these signature genes within the group of patients with a specific tumor type. The high and low designations could then be combined in an approach where a patient could be assigned as high or low in multiple groups. Survival analysis was conducted using the Cox proportional hazards model from the R package survival^25^, adjusting for confounding clinical factors such as tumor stage, gender and age. Subsequently, survival curves were derived using the Kaplan-Meier method within the same package. For visualization, the *ggsurvplot* function of the *survminer* package in R was utilized.

## References

1. Moretta, A., Bottino, C., Mingari, M.C., Biassoni, R. & Moretta, L. What is a natural killer cell? Nat Immunol 3 (2002).

2. Crinier, A. et al. High-Dimensional Single-Cell Analysis Identifies Organ-Specific Signatures and Conserved NK Cell Subsets in Humans and Mice. Immunity 49, 971–986.e975 (2018).

3. Cooper, M.A., Fehniger, T.A. & Caligiuri, M.A. The biology of human natural killer-cell subsets. Trends Immunol 22, 633–640 (2001).

4. Horowitz, A. et al. Genetic and environmental determinants of human NK cell diversity revealed by mass cytometry. Sci Transl Med 5, 208ra145 (2013).

5. Horowitz, A., et al. Class I HLA haplotypes form two schools that educate NK cells in different ways. Sci Immunol 1, eaag1672 (2016).

6. Goodridge, J.P., Önfelt, B. & Malmberg, K.-J. Newtonian cell interactions shape natural killer cell education. Immunol Rev 267, 197–213 (2015).

7. Björkström, N.K. et al. Expression patterns of NKG2A, KIR, and CD57 define a process of CD56dim NK-cell differentiation uncoupled from NK-cell education. Blood 116, 3853–3864 (2010).

8. Schlums, H. et al. Cytomegalovirus infection drives adaptive epigenetic diversification of NK cells with altered signaling and effector function. Immunity 42, 443–456 (2015).

9. Lopez-Vergès, S. et al. CD57 defines a functionally distinct population of mature NK cells in the human CD56dimCD16+ NK-cell subset. Blood 116, 3865–3874 (2010).

10. Juelke, K. et al. CD62L expression identifies a unique subset of polyfunctional CD56dim NK cells. Blood 116, 1299–1307 (2010).

11. Collins, P.L. et al. Gene Regulatory Programs Conferring Phenotypic Identities to Human NK Cells. Cell 176, 348–360.e312 (2019).

12. Smith, S.L. et al. Diversity of peripheral blood human NK cells identified by single-cell RNA sequencing. Blood Adv 4, 1388–1406 (2020).

13. Melsen, J.E. et al. Single-cell transcriptomics in bone marrow delineates CD56(dim)GranzymeK(+) subset as intermediate stage in NK cell differentiation. Frontiers in immunology 13, 1044398 (2022).

14. 14. Holmes, T.D., et al. The transcription factor Bcl11b promotes both canonical and adaptive NK cell differentiation. Sci Immunol 6, eabc9801 (2021).

15. Rückert, T., Lareau, C.A., Mashreghi, M.-F., Ludwig, L.S. & Romagnani, C. Clonal expansion and epigenetic inheritance of long-lasting NK cell memory. Nat Immunol 23, 1551–1563 (2022).

16. Tang, F. et al. A pan-cancer single-cell panorama of human natural killer cells. Cell 186, 4235–4251.e4220 (2023).

17. Rood, J.E., Maartens, A., Hupalowska, A., Teichmann, S.A. & Regev, A. Impact of the Human Cell Atlas on medicine. Nat Med 28, 2486–2496 (2022).

18. Yang, C. et al. Heterogeneity of human bone marrow and blood natural killer cells defined by single-cell transcriptome. Nat Commun 10 (2019).

19. Gayoso, A. et al. A Python library for probabilistic analysis of single-cell omics data. Nature Biotechnology 40, 163–166 (2022).

20. Haghverdi, L., Buettner, F. & Theis, F.J. Diffusion maps for high-dimensional single-cell analysis of differentiation data. Bioinformatics 31 (2015).

21. Aibar, S. et al. SCENIC: single-cell regulatory network inference and clustering. Nat Methods 14 (2017).

22. Scheiter, M. et al. Proteome Analysis of Distinct Developmental Stages of Human Natural Killer (NK) Cells. Molecular & Cellular Proteomics 12, 1099–1114 (2013).

23. Goodridge, J.P. et al. Remodeling of secretory lysosomes during education tunes functional potential in NK cells. Nat Commun 10, 514 (2019).

24. Xu, C. et al. Probabilistic harmonization and annotation of single-cell transcriptomics data with deep generative models. Mol Syst Biol 17 (2021).

25. Setty, M. et al. Characterization of cell fate probabilities in single-cell data with Palantir. Nature Biotechnology 37 (2019).

26. Bergen, V., Lange, M., Peidli, S., Wolf, F.A. & Theis, F.J. Generalizing RNA velocity to transient cell states through dynamical modeling. Nature Biotechnology (2020).

27. Manno, G.L. et al. RNA velocity of single cells. Nature 560, 494 (2018).

28. Wolf, F.A. et al. PAGA: graph abstraction reconciles clustering with trajectory inference through a topology preserving map of single cells. Genome Biology 20, 59 (2019).

29. Chaves, P. et al. Loss of Canonical Notch Signaling Affects Multiple Steps in NK Cell Development in Mice. J Immunol 201, 3307–3319 (2018).

30. Nagel, S. et al. Polycomb repressor complex 2 regulates HOXA9 and HOXA10, activating ID2 in NK/T-cell lines. Mol Cancer 9, 151 (2010).

31. Balzarolo, M., Watzl, C., Medema, J.P. & Wolkers, M.C. NAB2 and EGR-1 exert opposite roles in regulating TRAIL expression in human Natural Killer cells. Immunol Lett 151, 61–67 (2013).

32. Wiencke, J.K. et al. The DNA methylation profile of activated human natural killer cells. Epigenetics 11, 363–380 (2016).

33. Cho, Y. et al. The basic helix-loop-helix proteins differentiated embryo chondrocyte (DEC) 1 and DEC2 function as corepressors of retinoid X receptors. Mol Pharmacol 76, 1360–1369 (2009).

34. Adams, N.M. et al. Transcription Factor IRF8 Orchestrates the Adaptive Natural Killer Cell Response. Immunity 48, 1172–1182.e1176 (2018).

35. Mace, E.M. et al. Biallelic mutations in IRF8 impair human NK cell maturation and function. J Clin Invest 127, 306–320 (2017).

36. Wang, Y. et al. The IL-15-AKT-XBP1s signaling pathway contributes to effector functions and survival in human NK cells. Nat Immunol 20, 10–17 (2019).

37. Rabacal, W. et al. Transcription factor KLF2 regulates homeostatic NK cell proliferation and survival. Proc Natl Acad Sci U S A 113, 5370–5375 (2016).

38. Li, M. et al. DDIT3 Directs a Dual Mechanism to Balance Glycolysis and Oxidative Phosphorylation during Glutamine Deprivation. Adv Sci (Weinh) 8, e2003732 (2021).

39. Domínguez Conde, C., et al. Cross-tissue immune cell analysis reveals tissue-specific features in humans. Science 376, eabl5197 (2022).

40. Lotfollahi, M. et al. Mapping single-cell data to reference atlases by transfer learning. Nature Biotechnology, 1–10 (2021).

41. Combes, A.J., Samad, B. & Krummel, M.F. Defining and using immune archetypes to classify and treat cancer. Nat Rev Cancer 23, 491–505 (2023).

42. Dann, E., Henderson, N.C., Teichmann, S.A., Morgan, M.D. & Marioni, J.C. Differential abundance testing on single-cell data using k-nearest neighbor graphs. Nature Biotechnology 40, 245–253 (2022).

43. Nersesian, S. et al. NK cell infiltration is associated with improved overall survival in solid cancers: A systematic review and meta-analysis. Transl Oncol 14, 100930 (2021).

44. Cursons, J. et al. A Gene Signature Predicting Natural Killer Cell Infiltration and Improved Survival in Melanoma Patients. Cancer Immunol Res 7, 1162–1174 (2019).

45. Chu, T., Wang, Z., Pe’er, D. & Danko, C.G. Cell type and gene expression deconvolution with BayesPrism enables Bayesian integrative analysis across bulk and single-cell RNA sequencing in oncology. Nat Cancer 3, 505–517 (2022).

46. Colaprico, A. et al. TCGAbiolinks: an R/Bioconductor package for integrative analysis of TCGA data. Nucleic Acids Research 44, e71 (2016).

47. Goldman, M.J. et al. Visualizing and interpreting cancer genomics data via the Xena platform. Nature Biotechnology 38, 675–678 (2020).

48. Jin, S. et al. Inference and analysis of cell-cell communication using CellChat. Nat Commun 12, 1088 (2021).

49. Béziat, V., Descours, B., Parizot, C., Debré, P. & Vieillard, V. NK Cell Terminal Differentiation: Correlated Stepwise Decrease of NKG2A and Acquisition of KIRs. PLoS One 5, e11966 (2010).

50. Béziat, V. et al. NK cell responses to cytomegalovirus infection lead to stable imprints in the human KIR repertoire and involve activating KIRs. Blood 121, 2678–2688 (2013).

51. Lee, J. et al. Epigenetic modification and antibody-dependent expansion of memory-like NK cells in human cytomegalovirus-infected individuals. Immunity 42, 431–442 (2015).

52. Huang, D. et al. The neuronal chemokine CX3CL1/fractalkine selectively recruits NK cells that modify experimental autoimmune encephalomyelitis within the central nervous system. FASEB J 20, 896–905 (2006).

53. Cantoni, C. et al. NK Cells, Tumor Cell Transition, and Tumor Progression in Solid Malignancies: New Hints for NK-Based Immunotherapy? J Immunol Res 2016, 4684268 (2016).

54. Platonova, S. et al. Profound coordinated alterations of intratumoral NK cell phenotype and function in lung carcinoma. Cancer Res 71, 5412–5422 (2011).

55. Carrega, P. et al. CD56(bright)perforin(low) noncytotoxic human NK cells are abundant in both healthy and neoplastic solid tissues and recirculate to secondary lymphoid organs via afferent lymph. J Immunol 192, 3805–3815 (2014).

56. Lachota, M. et al. Mapping the chemotactic landscape in NK cells reveals subset-specific synergistic migratory responses to dual chemokine receptor ligation. EBioMedicine 96, 104811 (2023).

57. Brownlie, D. et al. Expansions of adaptive-like NK cells with a tissue-resident phenotype in human lung and blood. Proc Natl Acad Sci U S A 118, e2016580118 (2021).

58. Carrega, P. et al. Natural killer cells infiltrating human nonsmall-cell lung cancer are enriched in CD56 bright CD16(-) cells and display an impaired capability to kill tumor cells. Cancer 112, 863–875 (2008).

59. Rezaeifard, S., Talei, A., Shariat, M. & Erfani, N. Tumor infiltrating NK cell (TINK) subsets and functional molecules in patients with breast cancer. Mol Immunol 136, 161–167 (2021).

60. Bao, W. et al. HER2 interacts with CD44 to up-regulate CXCR4 via epigenetic silencing of microRNA-139 in gastric cancer cells. Gastroenterology 141, 2076–2087.e2076 (2011).

61. Xie, P. et al. CD44 potentiates hepatocellular carcinoma migration and extrahepatic metastases via the AKT/ERK signaling CXCR4 axis. Ann Transl Med 10, 689 (2022).

62. Zhu, X.S. et al. Transcriptional scaffold: CIITA interacts with NF-Y, RFX, and CREB to cause stereospecific regulation of the class II major histocompatibility complex promoter. Mol Cell Biol 20, 6051–6061 (2000).

63. Porse, B.T. et al. Loss of C/EBP alpha cell cycle control increases myeloid progenitor proliferation and transforms the neutrophil granulocyte lineage. J Exp Med 202, 85–96 (2005).

64. Colanzi, A. et al. Molecular mechanism and functional role of brefeldin A-mediated ADP-ribosylation of CtBP1/BARS. Proc Natl Acad Sci U S A 110, 9794–9799 (2013).

65. Bellelli, R. et al. POLE3-POLE4 Is a Histone H3-H4 Chaperone that Maintains Chromatin Integrity during DNA Replication. Mol Cell 72, 112–126.e115 (2018).

66. Li, S. et al. The transcription factors Egr2 and Egr3 are essential for the control of inflammation and antigen-induced proliferation of B and T cells. Immunity 37, 685–696 (2012).

## References

1. Bernstein, N.J. et al. Solo: Doublet Identification in Single-Cell RNA-Seq via Semi-Supervised Deep Learning. Cell Systems 11 (2020).

2. Wolf, F.A., Angerer, P. & Theis, F.J. SCANPY: large-scale single-cell gene expression data analysis. Genome Biology 19, 15 (2018).

3. Gayoso, A. et al. A Python library for probabilistic analysis of single-cell omics data. Nature Biotechnology 40, 163–166 (2022).

4. Luecken, M.D. et al. Benchmarking atlas-level data integration in single-cell genomics. Nat Methods 19, 41–50 (2022).

5. Wolf, F.A. et al. PAGA: graph abstraction reconciles clustering with trajectory inference through a topology preserving map of single cells. Genome Biology 20, 59 (2019).

6. Domínguez Conde, C., et al. Cross-tissue immune cell analysis reveals tissue-specific features in humans. Science 376, eabl5197 (2022).

7. Aibar, S. et al. SCENIC: single-cell regulatory network inference and clustering. Nat Methods 14 (2017).

8. Liberzon, A. et al. The Molecular Signatures Database (MSigDB) hallmark gene set collection. Cell systems 1 (2015).

9. Setty, M. et al. Characterization of cell fate probabilities in single-cell data with Palantir. Nature Biotechnology 37 (2019).

10. Manno, G.L. et al. RNA velocity of single cells. Nature 560, 494 (2018).

11. Bergen, V., Lange, M., Peidli, S., Wolf, F.A. & Theis, F.J. Generalizing RNA velocity to transient cell states through dynamical modeling. Nature Biotechnology (2020).

12. Van de Sande, B. et al. A scalable SCENIC workflow for single-cell gene regulatory network analysis. Nature Protocols 15 (2020).

13. Moerman, T. et al. GRNBoost2 and Arboreto: efficient and scalable inference of gene regulatory networks. Bioinformatics 35 (2019).

14. Islam, S. et al. Characterization of the single-cell transcriptional landscape by highly multiplex RNA-seq. Genome Res 21, 1160–1167 (2011).

15. Lotfollahi, M. et al. Mapping single-cell data to reference atlases by transfer learning. Nature Biotechnology, 1–10 (2021).

16. Jin, S. et al. Inference and analysis of cell-cell communication using CellChat. Nat Commun 12, 1088 (2021).

17. Murphy, A.E. & Skene, N.G. A balanced measure shows superior performance of pseudobulk methods in single-cell RNA-sequencing analysis. Nat Commun 13, 7851 (2022).

18. Robinson, M.D., McCarthy, D.J. & Smyth, G.K. edgeR: a Bioconductor package for differential expression analysis of digital gene expression data. Bioinformatics 26, 139–140 (2010).

19. Dann, E., Henderson, N.C., Teichmann, S.A., Morgan, M.D. & Marioni, J.C. Differential abundance testing on single-cell data using k-nearest neighbor graphs. Nature Biotechnology 40, 245–253 (2022).

20. Subramanian, A. et al. Gene set enrichment analysis: A knowledge-based approach for interpreting genome-wide expression profiles. Proceedings of the National Academy of Sciences 102, 15545–15550 (2005).

21. Colaprico, A. et al. TCGAbiolinks: an R/Bioconductor package for integrative analysis of TCGA data. Nucleic Acids Research 44, e71 (2016).

22. Goldman, M.J. et al. Visualizing and interpreting cancer genomics data via the Xena platform. Nature Biotechnology 38, 675–678 (2020).

23. Chu, T., Wang, Z., Pe’er, D. & Danko, C.G. Cell type and gene expression deconvolution with BayesPrism enables Bayesian integrative analysis across bulk and single-cell RNA sequencing in oncology. Nat Cancer 3, 505–517 (2022).

24. Tran, K.A. et al. Performance of tumour microenvironment deconvolution methods in breast cancer using single-cell simulated bulk mixtures. Nat Commun 14, 5758 (2023).

25. Therneau, T.M., Elizabeth, A. & Cynthia, C. survival: Survival Analysis. 2023.

