## Supplementary Figure and Tables for "Pan-cancer profiling of tumor-infiltrating natural killer cells through transcriptional reference mapping"

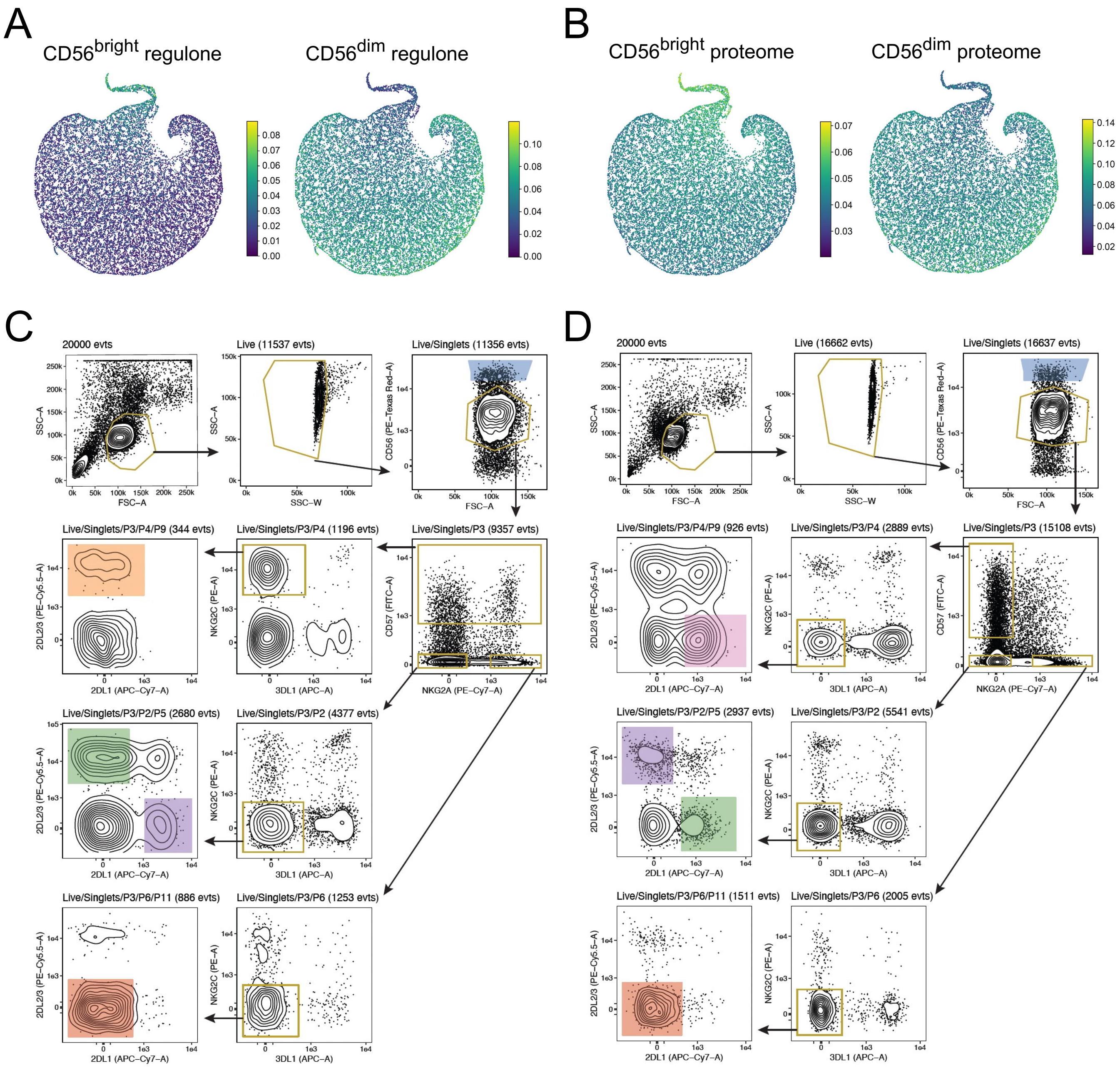


**Supplemental Figure 1.** Peripheral blood NK cell subsets and sorting strategy. (A-B) AUCell scores of gene signatures for CD56bright and CD56dim NK regulones (A) and proteomes (B). (C-D) Sorting strategy for phenotypically defined functional PB-NK cell subsets sequenced in one donor with (C) and one without (D) an adaptive expansion.

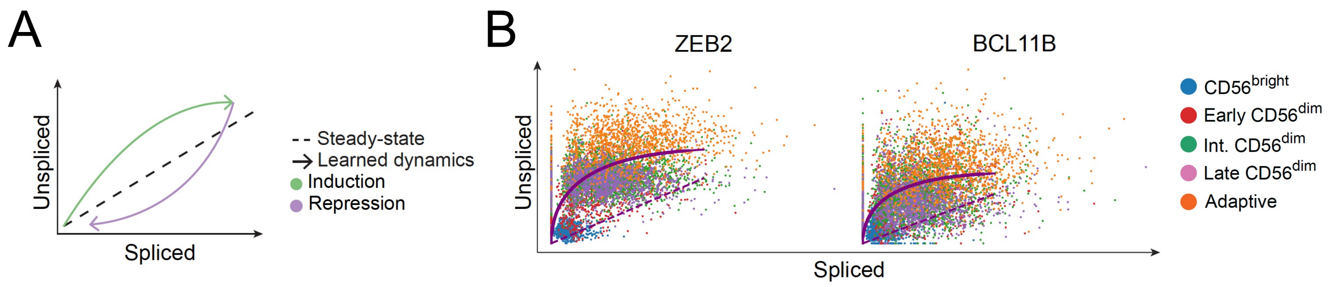


**Supplemental Figure 2.** RNA velocity. (A) Graphical depiction of inferring RNA velocity based on spliced vs unspliced transcripts. (B) RNA velocity plots for ZEB2 and BCL11B transcripts stratified by subset annotation in donors with an adaptive expansion.

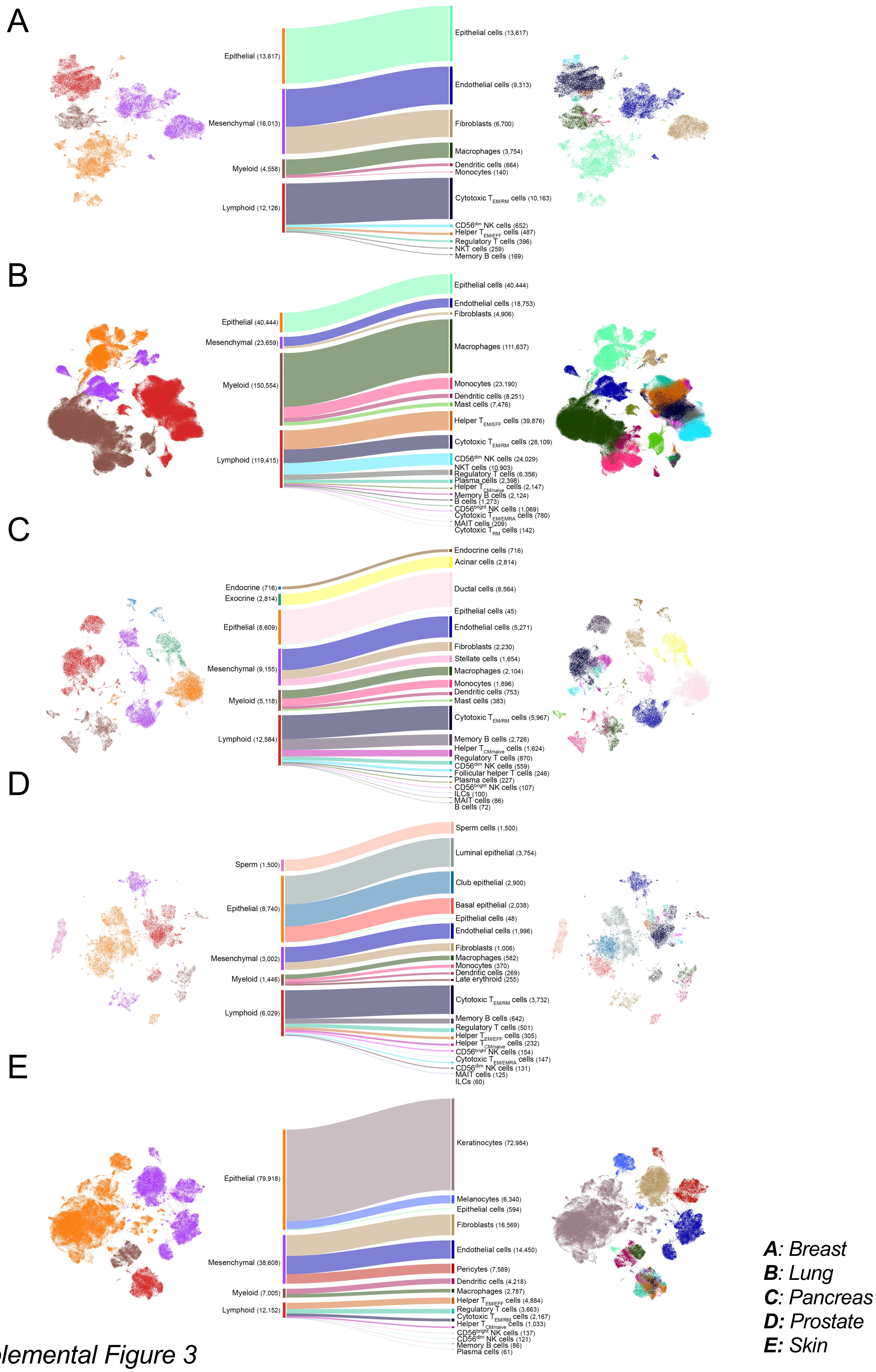


*Supplemental Figure 3*

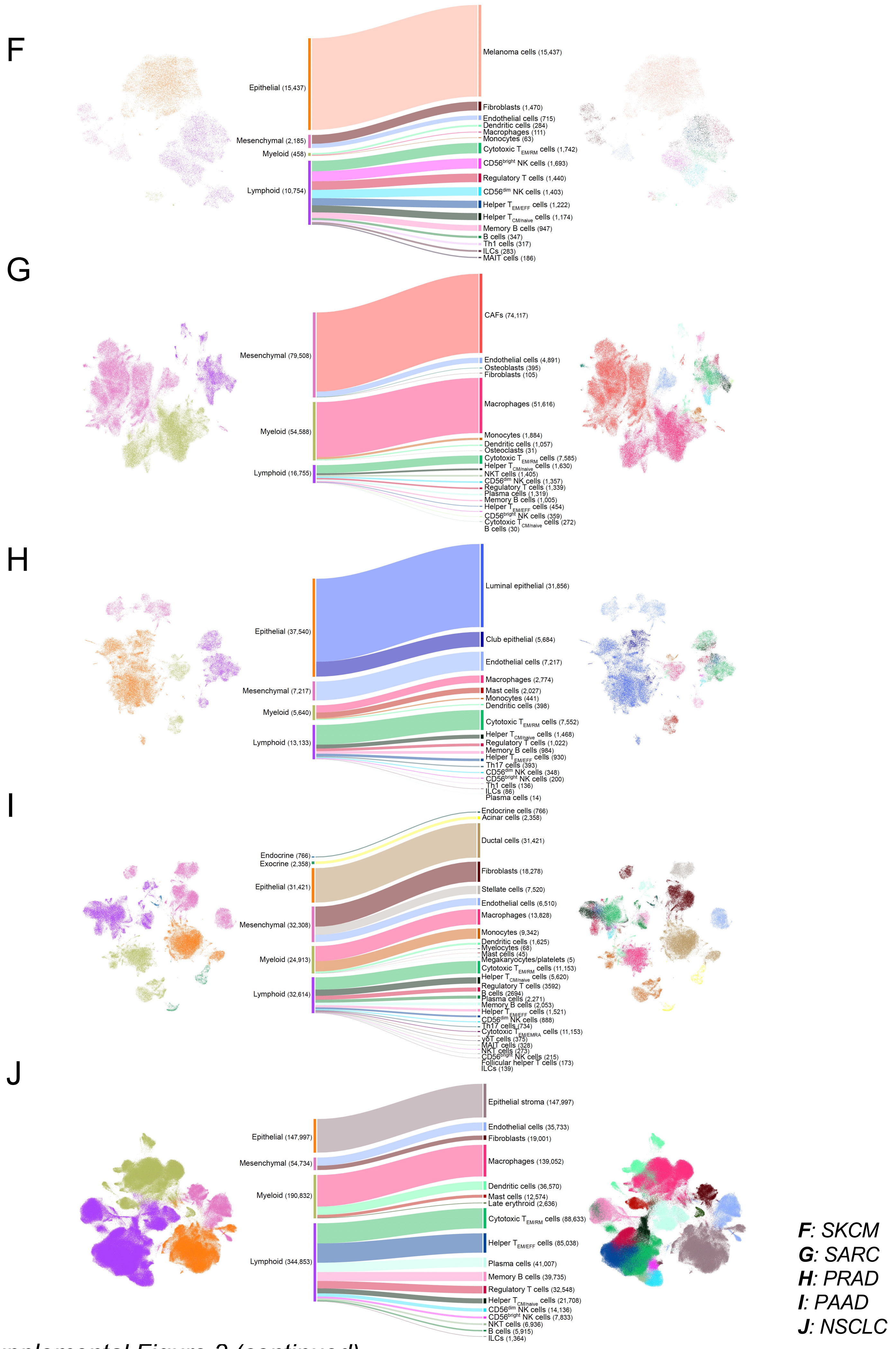


*Supplemental Figure 3 (continued)*

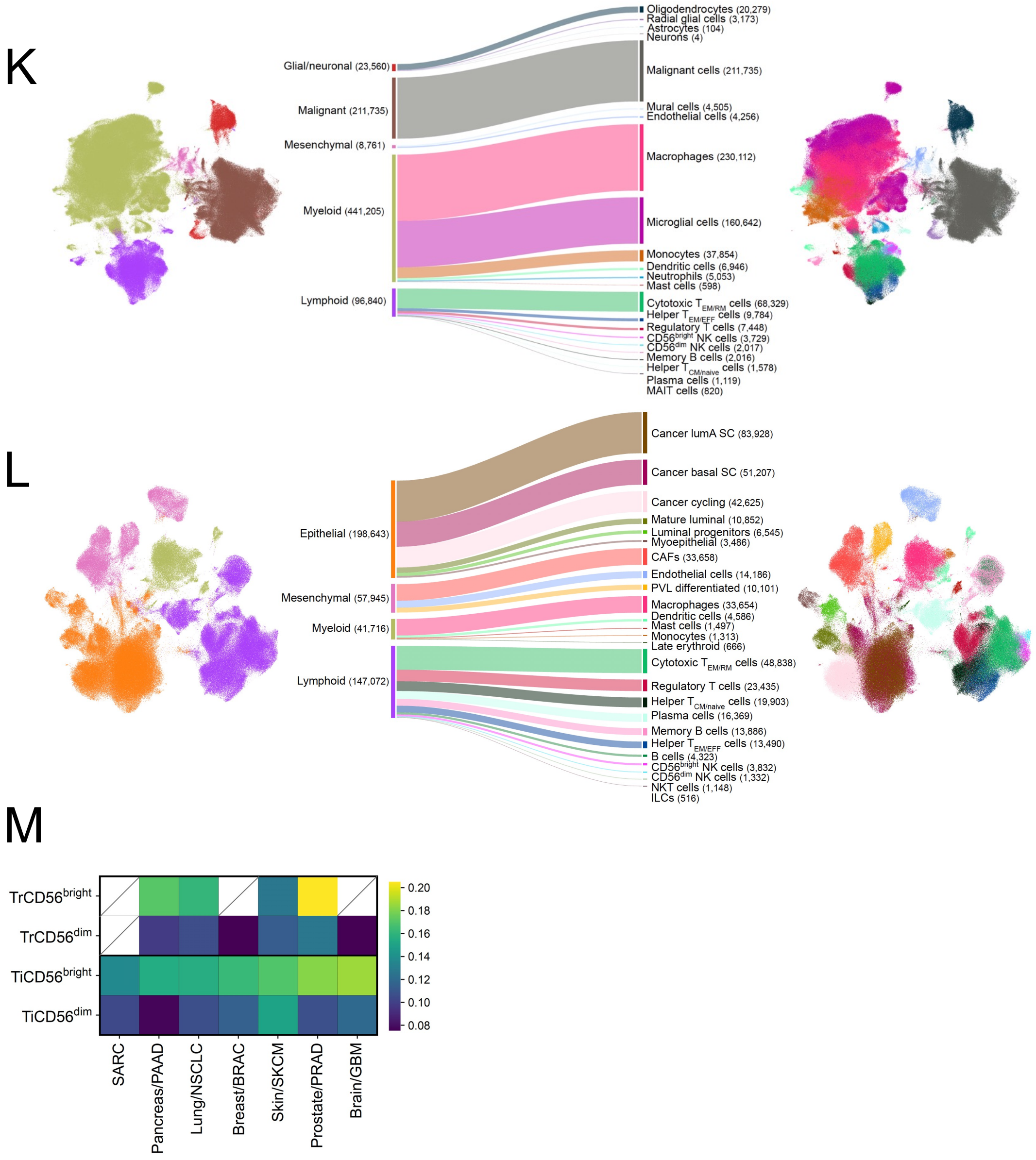


**Supplemental Figure 3.** Healthy tissue and solid-tumor dataset annotation using CellTypist. (A-L) Annotation of major cellular subtypes (left) which was further stratified into individual cell subtypes using CellTypist annotation (right) for Breast (A), Lung (B), Pancreas (C), Prostate (D), Skin (E), SKCM (F), SARC (G), PRAD (H), PAAD (I), NSCLC (J), GBM (K) and BRAC (L) datasets. (M) Tissue residency signature scoring for CD56bright and CD56dim NK cell subsets annotated from healthy tissue and solid-tumor datasets.

***K**: GBM*
***L**: BRAC*

*Supplemental Figure 3 (continued)*

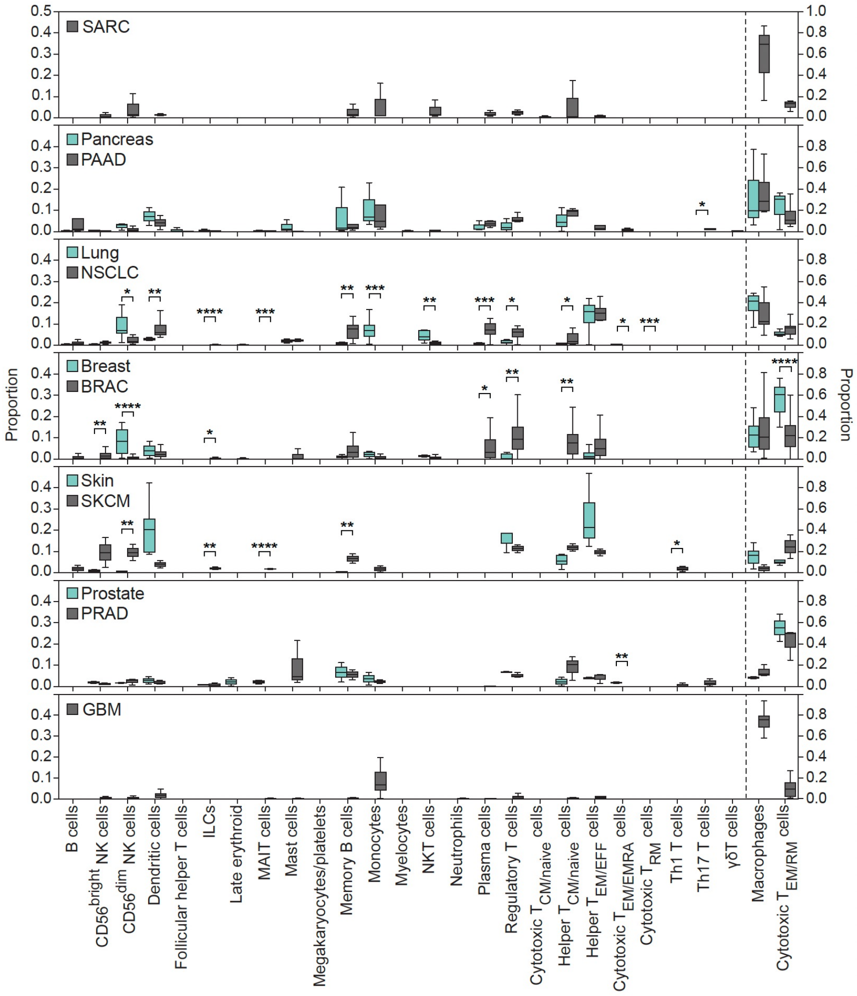


**Supplemental Figure 4.** Immune cell composition. Immune cell composition of healthy tissue and solid tumor datasets stratified by tumor type.

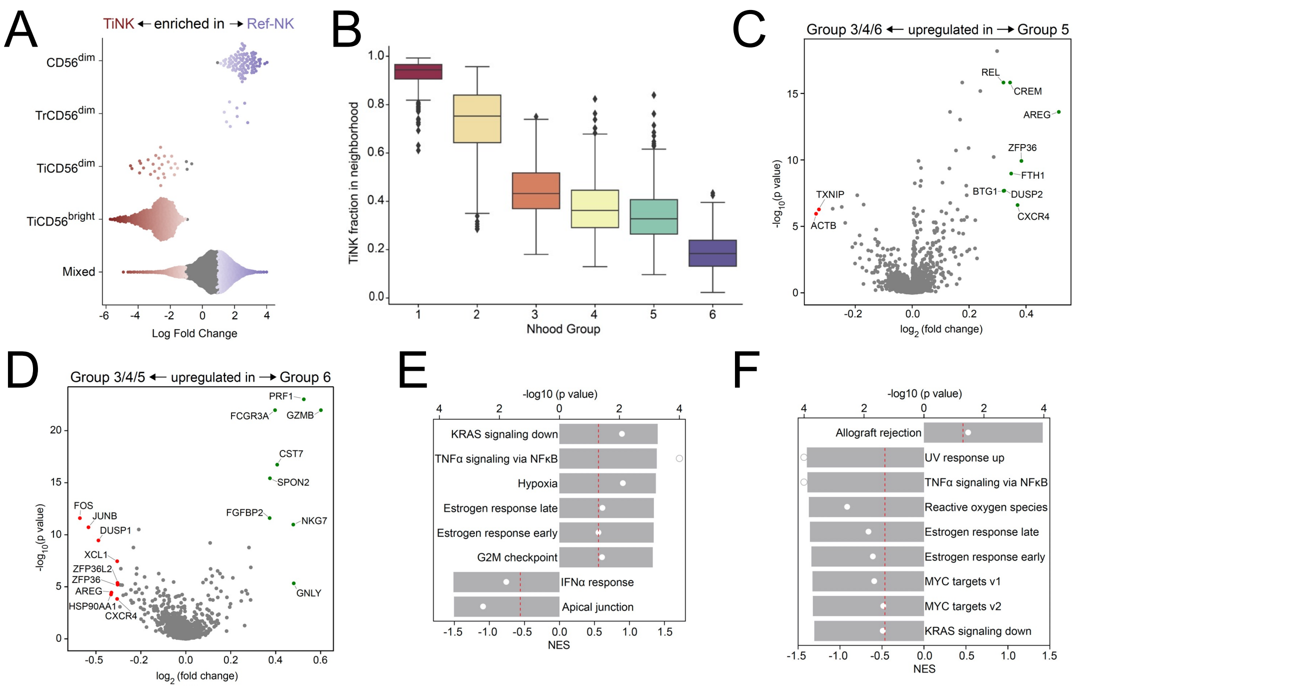


**Supplemental Figure 5.** Characterization of cellular states of NK cells identified in pan-cancer cell atlas. (A) Beaswarm plot depicting differential abundance of TiNK or Ref-NK (PB-NK, TrNK) enriched neighborhoods, clustered based on subset annotation of individual neighborhoods. (B) Fraction of TiNK cell specific neighborhoods within each neighborhood group. (C-F) Volcano plots depicting differentially expressed genes (DEGs) and corresponding gene set enrichment analysis (GSEA) between Group 5 vs. Group 3/4/6 (C, E) and Group 6 vs. Group 3/4/5 (D, F). Volcano plots: log fold change cutoff at 0.5, $p < 0.05$. GSEA plots: p value cutoff 0.5 (red line).

*Supplemental Figure 5*

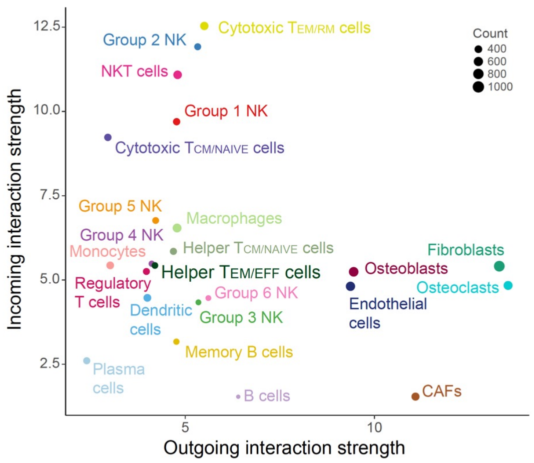


**Supplemental Figure 6.** Intercellular communication in SARC. Scatterplot depicting incoming and outgoing interaction strength of individual cell types in SARC as identified by CellChat.

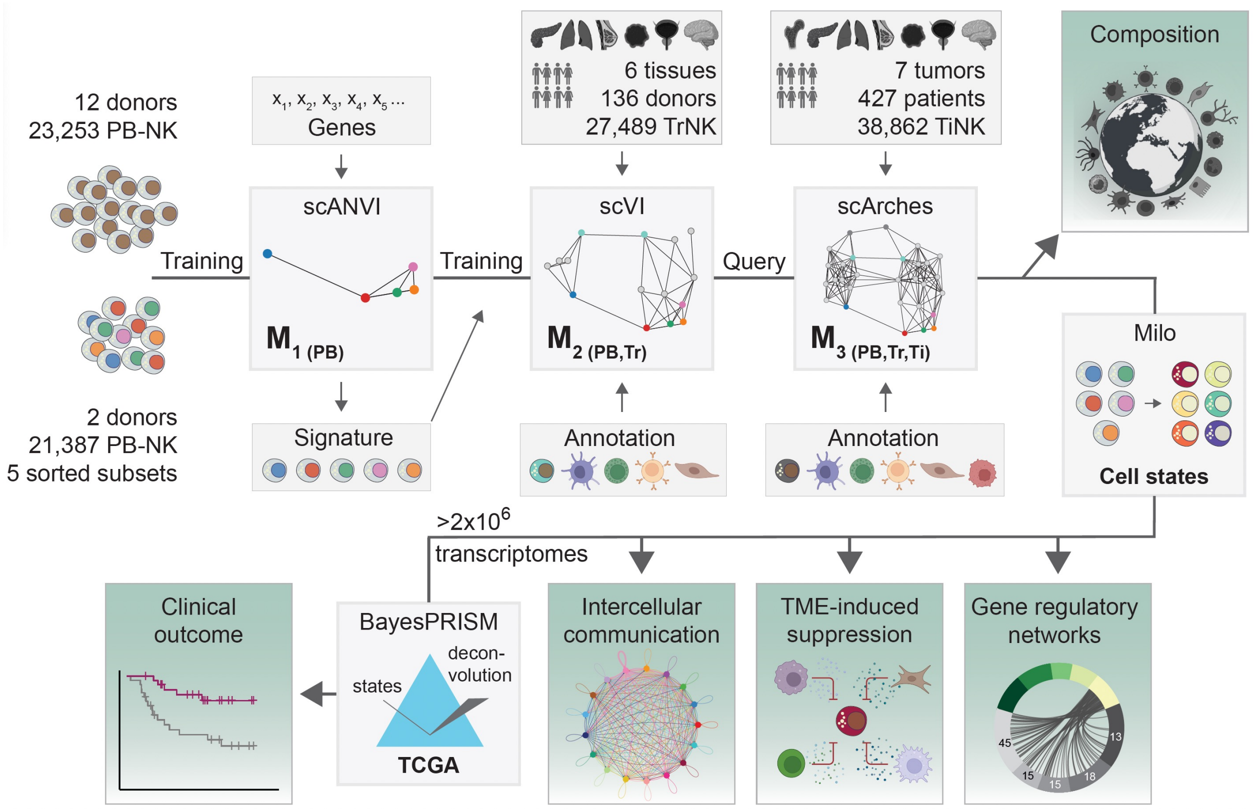


**Supplemental Figure 7.** Visualization of analysis workflow.

*Supplemental Figure 7*

### Supplemental tables

#### Supplemental table 1

Data containing NK cells from peripheral blood.

| Dataset | Number of donors | Platform | Comment | Citation |
|---|---|---|---|---|
| Bulk PB NK cells | 5 | 10x | Lab A in figure 1A | This manuscript |
| Sorted PB NK cells and bulk PB NK cells | 2 | 10x | Lab B in figure 1A | This manuscript |
| Bulk PB NK cells | 3 | 10x | Lab C in figure 1A | Crinier et al.[1] |
| Bulk PB NK cells | 2 | 10x | Lab D ini figure 1A | Yang et al.[2] |

#### Supplemental table 2

Data containing NK cells from normal tissue.

| Dataset | Tissue type | # patients | Platform | Comment | Citation |
|---|---|---|---|---|---|
| Bulk tissue | Lung | 7 | 10x | Tumor-adjacent normal lung samples, three from same patients as sub table 2 | Lambrechts et al.[3] |
| Bulk tissue | Lung | 4 | 10x | Tumor-adjacent normal lung samples, three from same patients as sub table 2 | Chan et al.[4] |
| Bulk tissue | Lung | 10 | 10x | Tumor-adjacent normal lung samples, same patients as sub table 2 | Bischoff et al.[5] |
| Bulk tissue | Lung | 7 | 10x | Tumor-adjacent normal lung samples, same patients as sub table 2 | Goveia et al.[6] |
| Bulk tissue | Lung | 11 | 10x | Tumor-adjacent normal lung samples, same patients as sub table 2 | Kim et al.[7] |
| Bulk tissue | Lung | 5 | 10x | Tumor-adjacent normal lung samples, same patients as sub table 2 | He et al.[8] |
| Bulk tissue | Lung | 42 | 10x | Tumor-adjacent normal lung samples, same patients as sub table 2 | Leader et al.[9] |
| Bulk tissue | Pancreas | 11 | 10x | 3 cases of non-pancreatic tumor patients (e.g. bile duct tumors or duodenal tumors) and 8 cases of non-malignant pancreatic tumor patients (e.g. pancreatic cyst) | Peng et al.[10] |
| Bulk tissue | Pancreas | 3 | 10x | adjacent/normal | Steele et al.[11] |
| Bulk tissue | Pancreas | 3 | 10x | adjacent/normal | Chen et al.[12] |
| Bulk tissue | Prostate | 10 | 10x | | Tuong et al.[13] |
| Bulk tissue | Prostate | 4 | 10x | | Heidegger et al.[14] |
| Bulk tissue | Breast | 10 | 10x | GEO: GSE161529 | Bhat et al.[15] |
| Bulk tissue | Brain | 4 | 10x | | Siletti et al.[16] |
| Bulk tissue | Skin | 3 | 10x | GEO: GSE169147 | Damskey et al.[17] |
| Bulk tissue | Skin | 6 | 10x | GEO: GSE193304 | He et al. |
| Bulk tissue | Skin | 3 | 10x | GEO: GSE160536 | Mirizio et al.[18] |
| Bulk tissue | Skin | 4 | 10x | GEO: GSE173205 | Rindler et al. |
| Bulk tissue | Skin | 4 | 10x | GEO: GSE138669 | Xue et al.[19] |

#### Supplemental table 3

Data containing NK cells from tumor tissue.

| Dataset | Tumor type | # patients | Platform | Comment | Accession | Citation |
|---|---|---|---|---|---|---|
| Bulk TME | Lung (NSCLC) | 8 | 10x | | ArrayExpress: E-MTAB-6149 (5 patients) and E-MTAB-6653 (3 patients) | Lambrechts et al.[3] |
| Bulk TME | Lung (NSCLC) | 7 | inDrop | | GEO: GSE127465 | Zilionis et al.[20] |
| Bulk TME | Lung (NSCLC) | 16 | 10x | HTAN | | Chan et al.[4] |
| Bulk TME | Lung (NSCLC) | 10 | 10x | | DOI: 10.24433/CO.0121060.v1 | Bischoff et al.[5] |
| Bulk TME | Lung (NSCLC) | 49 | 10x | scRNA-seq and CITE-seq | GEO: GSE154826 | Leader et al.[9] |
| Bulk TME | Lung (NSCLC) | 5 | 10x | | GSA: CRA001963 | He et al.[8] |
| Bulk TME | Lung (NSCLC) | 11 | 10x | | SRA: PRJNA634159 | Chen et al.[21] |
| Bulk TME | Lung (NSCLC) | 8 | 10x | | ArrayExpress: E-MTAB-6308 | Goveia et al.[6] |
| Bulk TME | Lung (NSCLC) | 11 | 10x | | GEO: GSE131907 | Kim et al.[7] |
| Bulk TME | Lung (NSCLC) | 42 | 10x | | GEO: GSE148071 | Wu et al.[22] |
| Bulk TME | Pancreas (PDAC) | 24 | 10x | | GSA: CRA001160 | Peng et al.[10] |
| Bulk TME | Pancreas (PDAC) | 10 | 10x | | GEO: GSE154778 | Lin et al.[23] |
| Bulk TME | Pancreas (PDAC) | 16 | 10x | | GEO: GSE155698 | Steele et al.[11] |
| Bulk TME | Pancreas (PDAC) | 6 | 10x | | GEO: GSE212966 | Chen et al.[12] |
| Bulk TME | Melanoma | 3 | 10x | Skin | GEO:GSE215120 | Zhang et al.[24] |
| Bulk TME | Melanoma | 8 | 10x | Skin | | Smalley et al.[25] |
| Bulk TME | Glioblastoma (GBM) | 4 | 10x | | GEO: GSE162631 | Xie et al.[26] |
| Bulk TME | Glioblastoma (GBM) | 7 | 10x | | Broad Insitute: SCP503 | Richards et al.[27] |
| Bulk TME | Glioblastoma (GBM) | 16 | 10x | | GEO: GSE182109 | Abdelfattah et al.[28] |
| Bulk TME | Glioblastoma (GBM) | 7 | 10x | | OSF: osf.io/4q32e/ | Ravi et al.[29] |
| Bulk TME | Glioblastoma (GBM) | 9 | 10x | | GEO: GSE131928 | Neftel et al.[30] |
| Bulk TME | Glioblastoma (GBM) | 3 | 10x | | GEO: GSE138794 | Wang et al.[31] |
| Bulk TME | Glioblastoma (GBM) | 3 | 10x | | GEO: GSE139448 | Wang et al.[32] |

| Dataset | Tumor type | # patients | Platform | Comment | Accession | Citation |
|---|---|---|---|---|---|---|
| Bulk TME | Glioblastoma (GBM) | 5 | 10x | | Synapse: syn22257780 | Johnson et al.[33] |
| Bulk TME | Glioblastoma (GBM) | 5 | 10x | | GEO: GSE163108 | Mathewson et al.[34] |
| Bulk TME | Glioblastoma (GBM) | 13 | 10x | | GEO: GSE163120 | Pombo et al.[35] |
| Bulk TME | Glioblastoma (GBM) | 35 | 10x | | GEO: GSE154795 | Lee et al.[36] |
| Bulk TME | Glioblastoma (GBM) | 10 | 10x | | GEO: GSE173278 | LeBlanc et al.[37] |
| Bulk TME | Breast cancer | 26 | 10x | | GEO: GSE176078 | Wu et al.[38] |
| Bulk TME | Breast cancer | 34 | 10x | | GEO: GSE161529 | Pal et al.[39] |
| Bulk TME | Breast cancer | 1 | GEXSCOPE | | GEO: GSE158399 | Xu et al.[40] |
| Bulk TME | Breast cancer | 5 | GEXSCOPE | | GEO: GSE180286 | Xu et al.[41] |
| Bulk TME | Breast cancer | 31 | 10x | treatment-naive pre-treatment | biokey. lambrechtslab.org | Bassez et al.[42] |
| Bulk TME | Prostate cancer | 4 | 10x | | GEO: GSE193337 | Heidegger et al.[14] |
| Bulk TME | Prostate cancer | 13 | 10x | | GEO: GSE141445 | Chen et al.[43] |
| Bulk TME | Prostate cancer | 10 | 10x | | | Tuong et al.[13] |
| Bulk TME | Sarcoma | 11 | 10x | Osteosarcoma | GEO: GSE152048 | Zhou et al.[44] |
| Bulk TME | Sarcoma | 6 | 10x | Osteosarcoma | GEO: GSE162454 | Liu et al.[45] |
| Bulk TME | Sarcoma | 4 | 10x | Osteosarcoma (TIL) | GEO: GSE198896 | Cillo et al. |

#### Supplemental table 4

Overview of number of cells from the various sources, including number of lymphoid cells and number of NK cells.

| Dataset | Total cells in dataset | Lymphoid cell | NK cells |
|---|---|---|---|
| Peripheral blood | | | 46303 |
| Lung tumor | 738416 | 350143 | 21887 |
| Breast tumor | 445376 | 149235 | 5075 |
| Glioblastoma | 782101 | 97438 | 5516 |
| Pancreas tumor | 124380 | 32659 | 1062 |
| Sarcoma | 150851 | 16316 | 1695 |
| Prostate | 66890 | 15171 | 515 |
| Melanoma | 28834 | 10754 | 1453 |
| Lung normal | 334072 | 119415 | 25098 |
| Breast normal | 46314 | 12126 | 652 |
| Pancreas normal | 38996 | 12584 | 657 |
| Prostate normal | 20717 | 6029 | 273 |
| Brain normal | | 3753 | 633 |
| Skin normal | 137683 | 12152 | 189 |
| ——- | ——— | ——— | ——— |
| Total | 2176214 | 837775 | 115500 |

#### References


1. Crinier, A. *et al.* High-Dimensional Single-Cell Analysis Identifies Organ-Specific Signatures and Conserved NK Cell Subsets in Humans and Mice. *Immunity* **49**, 971–986.e5 (2018).
2. Yang, C. *et al.* Heterogeneity of human bone marrow and blood natural killer cells defined by single-cell transcriptome. *Nat Commun* **10**, (2019).
3. Lambrechts, D. *et al.* Phenotype molding of stromal cells in the lung tumor microenvironment. *Nat Med* **24**, 1277–1289 (2018).
4. Chan, J. M. *et al.* Signatures of plasticity, metastasis, and immunosuppression in an atlas of human small cell lung cancer. *Cancer Cell* (2021) doi:10.1016/j.ccell.2021.09.008.
5. Bischoff, P. *et al.* Single-cell RNA sequencing reveals distinct tumor microenvironmental patterns in lung adenocarcinoma. *Oncogene* **40**, 6748–6758 (2021).
6. Goveia, J. *et al.* An Integrated Gene Expression Landscape Profiling Approach to Identify Lung Tumor Endothelial Cell Heterogeneity and Angiogenic Candidates. *Cancer Cell* **37**, 21–36.e13 (2020).
7. Kim, N. *et al.* Single-cell RNA sequencing demonstrates the molecular and cellular reprogramming of metastatic lung adenocarcinoma. *Nat Commun* **11**, 2285 (2020).
8. He, D. *et al.* Single-cell RNA sequencing reveals heterogeneous tumor and immune cell populations in early-stage lung adenocarcinomas harboring EGFR mutations. *Oncogene* **40**, 355–368 (2021).
9. Leader, A. M. *et al.* Single-cell analysis of human non-small cell lung cancer lesions refines tumor classification and patient stratification. *Cancer Cell* **39**, 1594–1609.e12 (2021).
10. Peng, J. *et al.* Single-cell RNA-seq highlights intra-tumoral heterogeneity and malignant progression in pancreatic ductal adenocarcinoma. *Cell Res* **29**, 725–738 (2019).
11. Steele, N. G. *et al.* Multimodal mapping of the tumor and peripheral blood immune landscape in human pancreatic cancer. *Nat Cancer* **1**, 1097–1112 (2020).
12. Chen, K. GEO: GSE212966. https://www.ncbi.nlm.nih.gov/geo/query/acc.cgi?acc=GSE212966.
13. Tuong, Z. K. *et al.* Resolving the immune landscape of human prostate at a single-cell level in health and cancer. *Cell Rep* **37**, 110132 (2021).
14. Heidegger, I. *et al.* Comprehensive characterization of the prostate tumor microenvironment identifies CXCR4/CXCL12 crosstalk as a novel antiangiogenic therapeutic target in prostate cancer. *Mol Cancer* **21**, 132 (2022).
15. Bhat-Nakshatri, P. *et al.* A single-cell atlas of the healthy breast tissues reveals clinically relevant clusters of breast epithelial cells. *Cell Rep Med* **2**, 100219 (2021).
16. Siletti, K. *et al.* Transcriptomic diversity of cell types across the adult human brain. 2022.10.12.511898 https://www.biorxiv.org/content/10.1101/2022.10.12.511898v1 (2022) doi:10.1101/2022.10.12.511898.
17. Damsky, W. *et al.* Inhibition of type 1 immunity with tofacitinib is associated with marked improvement in longstanding sarcoidosis. *Nat Commun* **13**, 3140 (2022).
18. Mirizio, E. *et al.* Single-cell transcriptome conservation in a comparative analysis of fresh and cryopreserved human skin tissue: Pilot in localized scleroderma. *Arthritis Res Ther* **22**, 263 (2020).
19. Xue, D. *et al.* Expansion of Fc$\gamma$ Receptor IIIa-Positive Macrophages, Ficolin 1-Positive Monocyte-Derived Dendritic Cells, and Plasmacytoid Dendritic Cells Associated With Severe Skin Disease in Systemic Sclerosis. *Arthritis Rheumatol* **74**, 329–341 (2022).
20. Zilionis, R. *et al.* Single-Cell Transcriptomics of Human and Mouse Lung Cancers Reveals Conserved Myeloid Populations across Individuals and Species. *Immunity* **50**, 1317–1334.e10 (2019).
21. Chen, J. *et al.* Single-cell transcriptome and antigen-immunoglobin analysis reveals the diversity of B cells in non-small cell lung cancer. *Genome Biol* **21**, 1–21 (2020).
22. Wu, F. *et al.* Single-cell profiling of tumor heterogeneity and the microenvironment in advanced non-small cell lung cancer. *Nat Commun* **12**, 2540 (2021).

23. Lin, W. *et al.* Single-cell transcriptome analysis of tumor and stromal compartments of pancreatic ductal adenocarcinoma primary tumors and metastatic lesions. *Genome Med* **12**, 1–14 (2020).

24. Zhang, C. *et al.* A single-cell analysis reveals tumor heterogeneity and immune environment of acral melanoma. *Nat Commun* **13**, 7250 (2022).

25. Smalley, I. *et al.* Single-Cell Characterization of the Immune Microenvironment of Melanoma Brain and Leptomeningeal Metastases. *Clin Cancer Res* **27**, 4109–4125 (2021).

26. Xie, Y. *et al.* Key molecular alterations in endothelial cells in human glioblastoma uncovered through single-cell RNA sequencing. *JCI Insight* **6**, e150861.

27. Richards, L. M. *et al.* Gradient of Developmental and Injury Response transcriptional states defines functional vulnerabilities underpinning glioblastoma heterogeneity. *Nat Cancer* **2**, 157–173 (2021).

28. Abdelfattah, N. *et al.* Single-cell analysis of human glioma and immune cells identifies S100A4 as an immunotherapy target. *Nat Commun* **13**, 767 (2022).

29. Ravi, V. M. *et al.* T-cell dysfunction in the glioblastoma microenvironment is mediated by myeloid cells releasing interleukin-10. *Nat Commun* **13**, 925 (2022).

30. Neftel, C. *et al.* An Integrative Model of Cellular States, Plasticity, and Genetics for Glioblastoma. *Cell* **178**, 835–849.e21 (2019).

31. Wang, L. *et al.* The Phenotypes of Proliferating Glioblastoma Cells Reside on a Single Axis of Variation. *Cancer Discov* **9**, 1708–1719 (2019).

32. Wang, R. *et al.* Adult Human Glioblastomas Harbor Radial Glia-like Cells. *Stem Cell Reports* **14**, 338–350 (2020).

33. Johnson, K. C. *et al.* Single-cell multimodal glioma analyses identify epigenetic regulators of cellular plasticity and environmental stress response. *Nat Genet* **53**, 1456–1468 (2021).

34. Mathewson, N. D. *et al.* Inhibitory CD161 receptor identified in glioma-infiltrating T cells by single-cell analysis. *Cell* **184**, 1281–1298.e26 (2021).

35. Pombo Antunes, A. R. *et al.* Single-cell profiling of myeloid cells in glioblastoma across species and disease stage reveals macrophage competition and specialization. *Nat Neurosci* **24**, 595–610 (2021).

36. Lee, A. H. *et al.* Neoadjuvant PD-1 blockade induces T cell and cDC1 activation but fails to overcome the immunosuppressive tumor associated macrophages in recurrent glioblastoma. *Nat Commun* **12**, 6938 (2021).

37. LeBlanc, V. G. *et al.* Single-cell landscapes of primary glioblastomas and matched explants and cell lines show variable retention of inter- and intratumor heterogeneity. *Cancer Cell* **40**, 379–392.e9 (2022).

38. Wu, S. Z. *et al.* A single-cell and spatially resolved atlas of human breast cancers. *Nature genetics* **53**, 1334 (2021).

39. Pal, B. *et al.* A single-cell RNA expression atlas of normal, preneoplastic and tumorigenic states in the human breast. *EMBO J* **40**, e107333 (2021).

40. Xu, K. *et al.* Integrative analyses of scRNA-seq and scATAC-seq reveal CXCL14 as a key regulator of lymph node metastasis in breast cancer. *Hum Mol Genet* **30**, 370–380 (2021).

41. Xu, K. *et al.* Single-cell RNA sequencing reveals cell heterogeneity and transcriptome profile of breast cancer lymph node metastasis. *Oncogenesis* **10**, 66 (2021).

42. Bassez, A. *et al.* A single-cell map of intratumoral changes during anti-PD1 treatment of patients with breast cancer. *Nat Med* **27**, 820–832 (2021).

43. Chen, S. *et al.* Single-cell analysis reveals transcriptomic remodellings in distinct cell types that contribute to human prostate cancer progression. *Nat Cell Biol* **23**, 87–98 (2021).

44. Zhou, Y. *et al.* Single-cell RNA landscape of intratumoral heterogeneity and immunosuppressive microenvironment in advanced osteosarcoma. *Nat Commun* **11**, 6322 (2020).

45. Liu, Y. *et al.* Single-Cell Transcriptomics Reveals the Complexity of the Tumor Microenvironment of Treatment-Naive Osteosarcoma. *Front Oncol* **11**, 709210 (2021).
